# Development of radiofluorinated MLN-4760 derivatives for PET imaging of the SARS-CoV-2 entry receptor ACE2

**DOI:** 10.1101/2024.03.20.585792

**Authors:** Jinling Wang, Darja Beyer, Christian Vaccarin, Yingfang He, Matthias Tanriver, Roger Benoit, Xavier Deupi, Linjing Mu, Jeffrey W. Bode, Roger Schibli, Cristina Müller

## Abstract

**Purpose:** The angiotensin converting enzyme 2 (ACE2) plays a regulatory role in the cardiovascular system and serves SARS-CoV-2 as an entry receptor. The aim of this study was to synthesize and evaluate radiofluorinated derivatives of the ACE2 inhibitor MLN-4760. [^18^F]F-MLN-4760 and [^18^F]F-Aza-MLN-4760 were demonstrated to be suitable for non-invasive imaging of ACE2, potentially enabling a better understanding of its expression dynamics.

**Methods:** Based on computational molecular modeling, the ACE2-binding modes of F-MLN-4760 and F-Aza-MLN-4760 were similar to that of MLN-4760. Co-crystallization of the hACE2/F-MLN-4760 protein complex was performed for confirmation. Displacement experiments using [^3^H]MLN-4760 enabled the determination of the binding affinities of the synthesized F-MLN-4760 and F-Aza-MLN-4760 to ACE2 expressed in HEK-ACE2 cells. Aryl trimethylstannane-based and pyridine-based radiofluorination precursors were synthesized and used for the preparation of the respective radiotracers. [^18^F]F-MLN-4760 and [^18^F]F-Aza-MLN-4760 were evaluated with regard to the uptake in HEK-ACE2 and HEK-ACE cells and in vitro binding to tissue sections of HEK-ACE2 xenografts and normal organs of mice. Biodistribution and PET/CT imaging studies of [^18^F]F-MLN-4760 and [^18^F]F-Aza-MLN-4760 were performed using HEK-ACE2 and HEK-ACE xenografted nude mice.

**Results:** Crystallography data revealed an equal ACE2-binding mode for F-MLN-4760 as previously found for MLN-4760 and indicated that the same would hold true for F-Aza-MLN-4760. The IC_50_ values were all in the high nM range, but three-fold lower for F-MLN-4760 and seven-fold lower for F-Aza-MLN-4760 than for MLN-4760. [^18^F]F-MLN-4760 and [^18^F]F-Aza-MLN-4760 were obtained in 1.4 ± 0.3 GBq and 0.5 ± 0.1 GBq activity with >99% radiochemical purity in a 5.3% and 1.2% radiochemical yield, respectively. Uptake in HEK-ACE2 cells was higher for [^18^F]F-MLN-4760 (67 ± 9%) than for [^18^F]F-Aza-MLN-4760 (37 ± 8%) after 3 h incubation while negligible uptake was seen in HEK-ACE cells (<0.3%). [^18^F]F-MLN-4760 and [^18^F]F-Aza-MLN-4760 accumulated specifically in HEK-ACE2 xenografts of mice (13 ± 2% IA/g and 15 ± 2% IA/g at 1 h p.i.) with almost no uptake observed in HEK-ACE xenografts (<0.3% IA/g). This was confirmed by PET/CT imaging, which also visualized unspecific accumulation in the gall bladder and intestinal tract.

**Conclusion:** Both radiotracers showed specific and selective binding to ACE2 in vitro and in vivo. [^18^F]F-MLN-4760 was, however, obtained in higher yields and the ACE2-binding affinity was superior over that of [^18^F]F-Aza-MLN-4760. [^18^F]F-MLN-4760 would, thus, be the candidate of choice for further developlment to enable PET imaging of ACE2 in patients.

## Introduction

In March 2020, the World Health Organization announced the coronavirus disease 2019 (Covid-19) pandemic due to the worldwide increasing number of infections by the acute severe respiratory syndrome Coronavirus-2 (SARS-CoV-2) and the poorly understood risk factors for severe disease progression [1, 2]. It is known that SARS-CoV-2 interacts with the human angiotensin converting enzyme 2 (hACE2) and uses it as a cell entry receptor [3]. ACE2 is an enzyme involved in the renin-angiotensin-aldosterone-system (RAAS), as is the case also for the better-known angiotensin converting enzyme (ACE) [4]. The primary functions of ACE are the conversion of angiotensin I to angiotensin II and the hydrolysis of bradykinin, which leads to vasoconstriction and related effects. ACE2 plays a counterregulatory role by cleaving angiotensin II to angiotensin 1-7 which interacts with the MAS receptor to induce vasodilatation and, hence, cardioprotective, anti-inflammatory effects. The interplay of ACE and ACE2 is essential for the regulation of the cardiovascular system (Fig. 1)

**Fig. 1.**
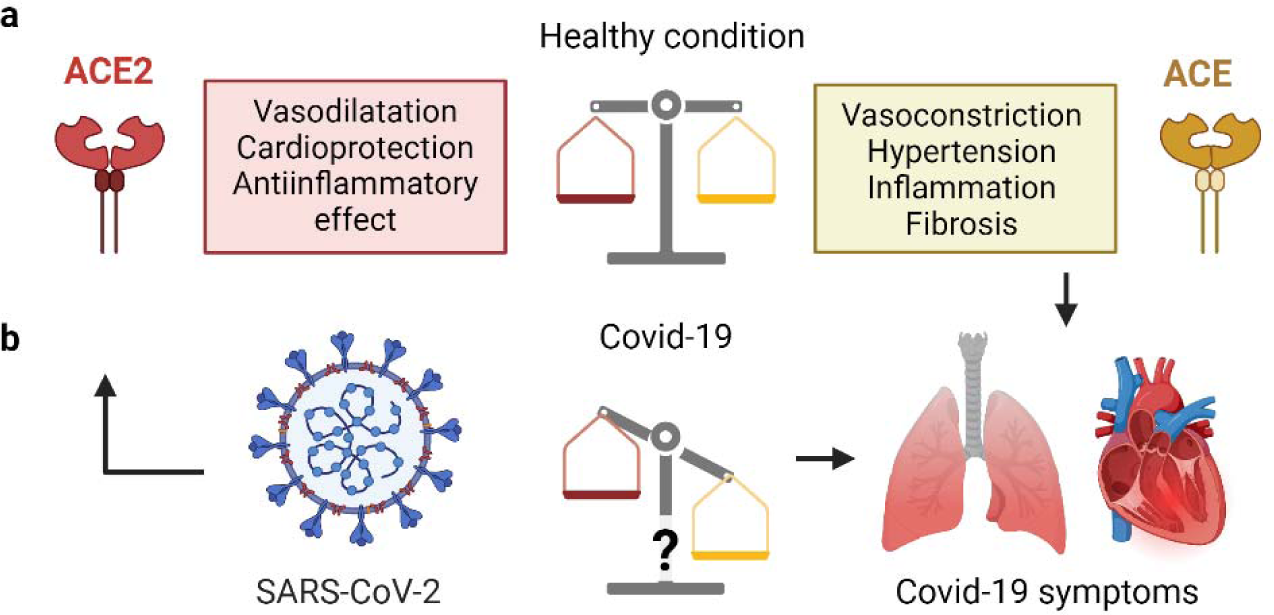
**a** Schematic representation of the counterregulatory enzymes, ACE and ACE2, which are relevant for the renin-angiotensin-aldosterone system (RAAS); **b** ACE2 serves as an entry receptor for SARS-CoV-2; as a result of the viral infection the RAAS may be affected.

It remains unclear to which extent the expression level and dynamics of ACE2 contribute to variations in infection susceptibility among individuals and whether the function of ACE2 is affected upon infection by SARS-CoV-2 [5, 6]. A hypothetical downregulation of ACE2 upon SARS-CoV-2 infection may lead to increased angiotensin II levels and related symptoms, which are consistent with clinical symptoms observed in severe Covid-19 cases [7–9]. This is in line with the fact that alteration of ACE2 expression levels were associated with cardiac and vascular pathophysiologies such as hypertension, ischemia and atherosclerosis [10–12].

Non-invasive imaging of ACE2 may contribute to a better understanding of the regulation of this important enzyme and potentially identify patients at risk for severe Covid-19 disease outcomes. Among the various imaging technologies, positron emission tomography (PET) is particularly useful to investigate (patho)physiological processes in cardiovascular diseases [13]. Several PET imaging studies have been performed at many nuclear medicine centers worldwide using [^18^F]fluorodeoxyglucose ([^18^F]FDG) to detect inflammation and pulmonary lesions in patients with Covid-19 [14–16]. Target-specific radiotracers for imaging ACE2 were also proposed [17]. Several preclinical studies reported the development of DX600, a ACE2-targeting cyclic peptide, derivatized with a DOTA or NODAGA chelator to allow the labeling with gallium-68, copper-64 or aluminium-fluoride-18 [18–20]. The Al[^18^F]F-NOTA-DX600 radiopeptide is currently being investigated in a clinical trial for non-invasive mapping of ACE2 (NCT04542863) [21]. Using this same concept, we derivatized DX600 also with an HBED-CC chelator for labeling with gallium-67/68 [22]. [^67^Ga]Ga-HBED-CC-DX600 showed distinct uptake in HEK-ACE2 xenografts and rapid clearance from the blood and kidneys. Despite these promising data, the relatively low hACE2-binding affinity of DX600-based radiopeptides (K_D_ ∼100 nM) may not be sufficient for imaging low expression levels of ACE2 in patients. This could explain the low uptake of the DX600-based [^68^Ga]Ga-HZ20 in the lung and heart of patients [20].

The aim of this study was to develop a ^18^F-based radiotracer for PET imaging of ACE2. The small molecular weight ACE2 inhibitor MLN-4760 was identified as a suitable lead structure as it was reported to bind to hACE2 with high affinity (IC_50_ = 0.44 nM [23]) and to show cross reactivity to the murine ACE2 but no binding to ACE (Fig. 2a) [24, 25]. The radiosynthesis of [^18^F]F-MLN-4760 and [^18^F]F-Aza-MLN-4760 was established based on specifically designed precursor molecules (Fig. 2 b/c). The non-radioactive reference compounds, F-MLN-4760 and F-Aza-MLN-4760, were synthesized to determine ACE2 binding affinity. The radiotracers were evaluated using HEK-ACE2 and HEK-ACE cells as well as the respective xenograft mouse model which was previously established in the group [22].

**Fig. 2.**
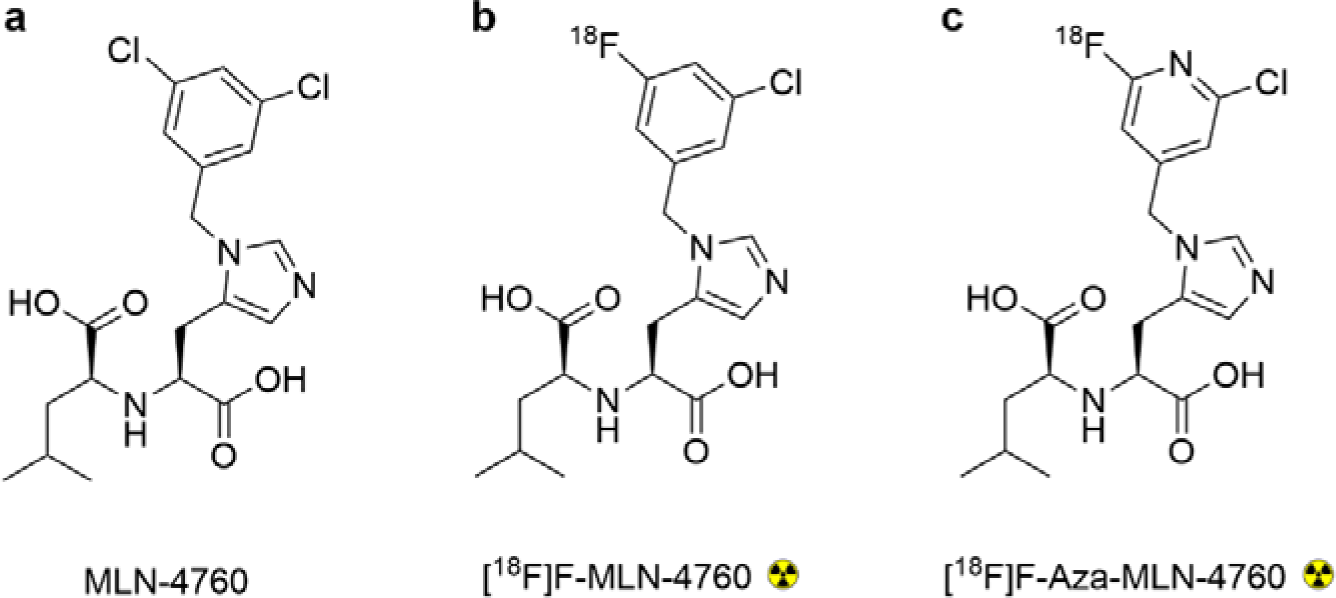
**a** Chemical structures of lead structure MLN-4760; **b/c** Chemical structures of the PET radiotracers, [^18^F]F-MLN-4760 and [^18^F]F-Aza-MLN-4760, to be developed and investigated.

## Methods

### Computational modeling and co-crystallization of hACE2-bound compounds

Computational molecular modeling techniques were applied to predict the binding mode of F-MLN-4760 and F-Aza-MLN-4760 in hACE2, based on a previously reported crystal structure of the hACE2/MLN-4760 complex (Supplementary Material; Fig. S1) [26].

The extracellular part of hACE2 with a C-terminal His-tag was expressed in insect cells and purified to achieve a quality corresponding to crystallization standard (Supplementary Material). Crystals of the concentrated protein, complexed with F-MLN-4760, were grown. A dataset to 2.5 Å was collected from a single crystal at 100 K at the X06SA beamline of the Swiss Light Source synchrotron at the Paul Scherrer Institute, using a wavelength of 1.0 Å. The data was processed using XDS X-ray detector software [27]. The structure was solved by molecular replacement with Phaser [28] using an AlphaFold model [29] of hACE2 as the search model. There were two hACE2 molecules in the asymmetric unit. The model was built and refined iteratively [30, 31] and the geometry and stereochemistry were validated [32].

### Synthesis of MLN-4760, F-MLN-4760 and F-Aza-MLN-4760

MLN-4760 was synthesized according to previously published procedures with slight modifications (Supplementary Material; Scheme S1) [23, 33]. F-MLN-4760 was prepared according to the same synthesis approach as applied for MLN-4760, however, instead of using (3,5-dichlorophenyl)methanol the (3-chloro-5-fluorophenyl)methanol was employed in the respective synthesis step (Supplementary Material; Scheme S1). F-Aza-MLN-4760 was produced based on a novel synthesis approach that involved *tert*-butyl protection of the carboxylic groups (Supplementary Material; Scheme S2). HPLC purification of the (*S,S*)-MLN-4760 was based on the comparison with the commercially available compound (MLN-4760; MW: 428.31; Merck, CAS N° 305335-31-3). The isolation of the desired (*S*,*S*)-diastereoisomers of the respective fluorinated MLN-4760 derivatives was based on the assumption that the HPLC elution sequence of the diastereoisomers would be identical to that of the MLN-4760 lead compound.

### Synthesis of precursor molecules for the preparation of the radiotracers

The synthetic approach adopted for the preparation of the aryl-trimethylstannane precursor **1** was based on the same reaction sequence described for F-Aza-MLN-4760 (Supplementary Material; Scheme S2). This allowed us to obtain a *tert*-butyl-protected compound suitable for the subsequent radiolabeling step. One of the chlorine atoms on the aromatic ring of MLN-4760 was substituted with iodine to enable a palladium-catalyzed stannylation as the last synthetic step (Supplementary Material, Scheme S3). The production of the pyridine-based precursor **2** was based on the same reaction sequence, however, a chlorinated pyridine was employed as an alkylating agent in the respective synthesis step (Supplementary Material, Scheme S4). The desired (*S*,*S*)-diastereomers of both precursor **1** and **2** were isolated by preparative HPLC based on the assumption that the elution sequence of the diastereoisomers would be identical to that of the MLN-4760 lead compound.

### Radiosynthesis of [^18^F]F-MLN-4760 and [^18^F]F-Aza-MLN-4760

Precursors **1** and **2** were used for ^18^F-fluorination to prepare [^18^F]F-MLN-4760 and [^18^F]F-Aza-MLN-4760, respectively (Scheme 1). The synthesis of [^18^F]F-MLN-4760 was performed by mixing the precursor **1**, Cu(OTf)_2_(Py)_4_ and azeotropically dried K[^18^F]F in *N,N*-dimethylacetamide followed by heating at 110 °C for 10 min. The subsequent deprotection reaction was performed by adding 85% orthophosphoric acid and heating at 110 °C for 15 min (Supplementary Material). [^18^F]F-Aza-MLN-4760 was prepared by heating a solution of precursor **2**, Kryptofix^®^ 222 and azeotropically dried Cs[^18^F]F in dimethylsulfoxide at 195 °C for 20 min. The ^t^Bu groups were cleaved by the addition of HCl 4 M and further stirring at 80 °C for 20 min (Supplementary Material). After partial neutralization of the acidic reaction mixtures, both [^18^F]F-MLN-4760 and [^18^F]F-MLN-4760 were purified by semipreparative HPLC and subsequently trapped using a C18 or cation-exchange cartridge, respectively, which allowed the formulation of these radiotracers in phosphate buffered saline (PBS) using a maximum ethanol content of 10% (*v/v*).

**Scheme 1.**
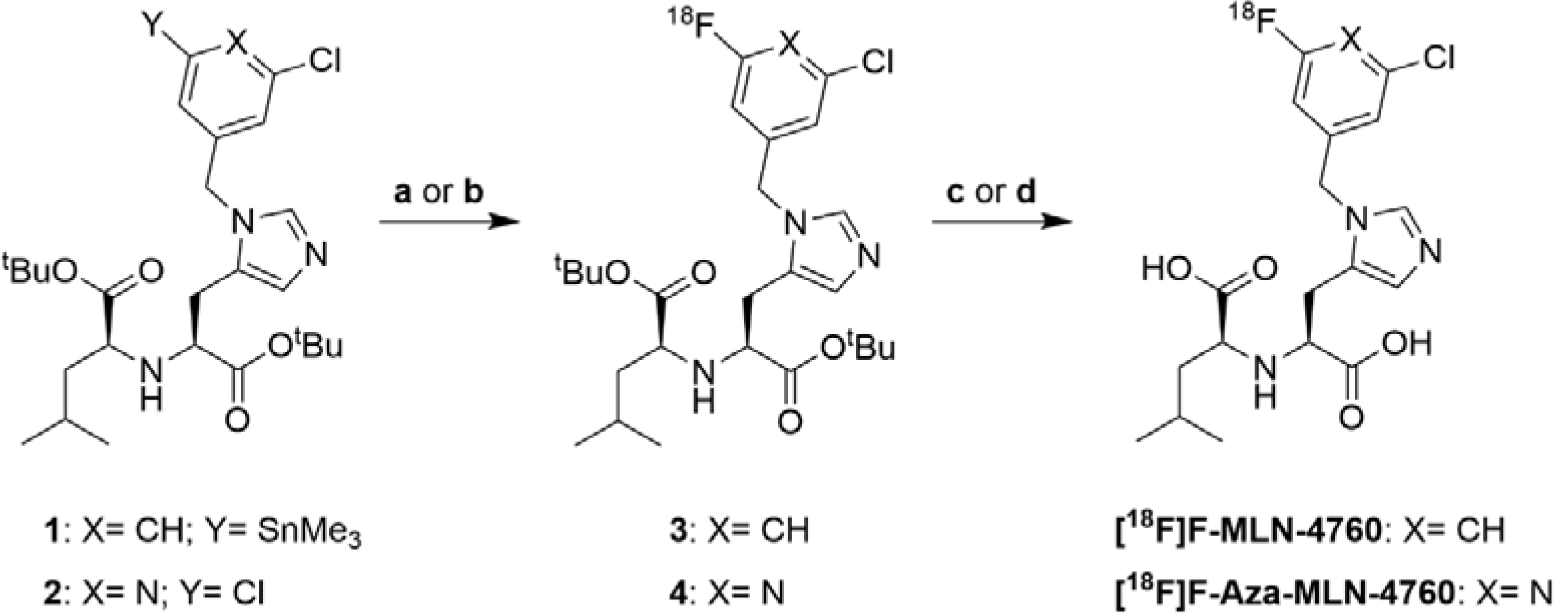
Radiofluorination of the precursors to obtain [^18^F]F-MLN-4760 and [^18^F]F-Aza-MLN-4760. Reaction conditions: (**a**) K[^18^F]F, Cu(OTf)_2_(Py)_4_, DMA, 110 °C, 10 min; (**b**) Cs[^18^F]F, DMSO, 195 °C, 20 min; (**c**) H_3_PO_4_ (85%, *m/m*), 110 °C, 15 min; (**d**) HCl (4 M), 80 °C, 20 min.

### In vitro stability and distribution coefficient of [^18^F]F-MLN-4760 and [^18^F]F-Aza-MLN-4760

The radiolytic stability of [^18^F]F-MLN-4760 and [^18^F]F-Aza-MLN-4760 formulated in PBS pH 7.4 containing 10% EtOH was investigated over 3 h incubation at RT (Supplementary Material). The metabolic stability was determined after incubation of the radiotracers in murine and human blood plasma at 37 °C (Supplementary Material). A shake flask method was employed to determine the *n*-octanol/PBS pH 7.4 distribution coefficient (logD value) of the ^18^F-based radiotracers as previously reported (Supplementary Material) [22]. The result was presented as the average value (± standard deviation, SD) of three independent logD experiments each performed with five replicates.

### Cell culture

HEK cells transfected with hACE2 (HEK-ACE2) or hACE (HEK-ACE), respectively, were obtained from Innoprot (Innovative Technologies in Biological Systems S.L. Bizkaia, Spain) and cultured in Dulbecco’s Modified Eagle Medium (DMEM) supplemented with non-essential amino acids, fetal calf serum, antibiotics and hygromycin B using standard culture conditions (Supplementary Material) [22].

### ACE2-binding affinity of F-MLN-4760 and F-Aza-MLN-4760

The IC_50_-values of MLN-4760, F-MLN-4760 and F-Aza-MLN-4760 were determined using HEK-ACE2 cells (Supplementary Material). In brief, the cells were seeded in 48-well plates, allowing adhesion and growth overnight. [^3^H]MLN-4760 (RC Tritec AG, Teufen, Switzerland) was added to each well (10 µL, 3.2 pmol, 6.2 nM) together with increasing concentrations (100 µM - 0.01 nM) of the test agents, MLN-4760, F-MLN-4760 or F-Aza-F-MLN-4760. After a 1 h incubation time at 37 °C, the cells were rinsed with PBS followed by lysis using NaOH (1 M, 600 µL) and transferred to scintillation vials. After the addition of 5 mL scintillation cocktail (Ultima Gold, Perkin Elmer^®^), the samples were counted in a scintillation counter (TRI-CARB® 2250CA, Packard) and the counts were plotted against the logarithmic concentration of the test agents to obtain the IC_50_ values using GraphPad Prism software (version 8.3.1). The relative binding affinities were expressed as the reversed IC_50_ values which were standardized to MLN-4760 set as 1.0.

### Uptake of ^18^F-radiotracers in HEK-ACE2 and HEK-ACE cells

The uptake and internalization of [^18^F]F-MLN-4760 and [^18^F]F-Aza-MLN-4760 in HEK-ACE2 and HEK-ACE cells were investigated according to a previously published procedure (Supplementary Material)[22]. In brief, cells were seeded in 12-well plates and let to grow and adhere overnight. After the addition of the radiotracer, the cells were incubated for 1 h or 3 h in fresh medium. The supernatant was removed and the cells were rinsed with PBS or additionally with acidic stripping buffer to determine the total uptake and the internalized fraction, respectively. The cells were lysed with NaOH (1 M, 1 mL) and transferred to RIA tubes for counting in a γ-counter (Perkin Elmer, Wallac Wizard 1480). The uptake and internalized fraction of the radiotracers were expressed as the percentage of total added activity to each well.

### In vitro autoradiography studies

Autoradiography studies were performed on 10-µm thick frozen tissue sections of kidneys, heart, brain and lung tissue of CD1 nude mice as well as HEK-ACE and HEK-ACE2 xenografts. The tissue sections were thawed and hydrated in Tris-HCl buffer containing 0.25% BSA followed by incubation with [^18^F]F-MLN-4760 or [^18^F]F-Aza-MLN-4760 (225 kBq/150 µL) in Tris-HCl buffer containing 1% BSA with or without excess of MLN-4760 (10 µM) for 1 h at room temperature. After several rinsing steps and drying of the tissue sections, autoradiographic images were obtained after exposure of the tissue sections to a phosphor screen (Super resolution using a storage phosphor system Cyclone Plus; Perkin Elmer). Quantification of the signals was performed using OptiQuant software (version 5.0) based on a standards exposed at the same time. The unspecific binding of the radiotracers was determined in the presence of excess MLN-4760, which blocked the binding sites of ACE2. This value was subtracted from the total binding to obtain the specific binding of the radiotracers on each tissue section. The presented data were obtained from two independent experiments performed in duplicates of normal organs and xenografts from 2-3 different mice.

### In vivo studies

All applicable international, national, and/or institutional guidelines for the care and use of animals were followed. In particular, all animal experiments were carried out according to the guidelines of the Swiss Regulations for Animal Welfare. The preclinical studies have been ethically approved by the Cantonal Committee of Animal Experimentation and permitted by the responsible cantonal authorities (License N° 75743).

### Biodistribution and PET/CT imaging studies

Five-week-old female Crl:CD1-*Foxn^nu^* (CD1 nude) mice were obtained from Charles River Laboratories (Sulzfeld, Germany). Subcutaneous inoculation of HEK-ACE2 cells (8 × 10^6^ cells in 100 µL PBS pH 7.4) on the right shoulder and HEK-ACE cells (4 ×10^6^ cells in 100 µL PBS pH 7.4) on the left shoulder was performed as previously reported [22]. Biodistribution studies were performed 2-4 weeks later by intravenous injection of [^18^F]F-MLN-4760 or [^18^F]F-Aza-MLN-4760 (5 MBq, 100 µL) diluted in NaCl 0.9% containing 0.05% bovine serum albumin (BSA). Mice were sacrificed and dissected at 15 min, 1 h or 3 h after injection. Selected organs and tissues were collected, weighed and counted in a γ-counter for activity at the same time as standards of the original injection solution. The decay-corrected data were expressed as percent of the injected activity per gram of tissue mass (% IA/g). PET/CT imaging was performed at 15 min, 1 h and 3 h after injection of the radiotracers using a small-animal PET/CT scanner (G8 PET/CT; SOFIE, Dulles, U.S.A. [34]) as previously reported (Supplementary Material) [35].

## Results

### Computational model and crystal structure of hACE2 complexes of MLN-4760 derivatives

Computational models of F-MLN-4760 and F-Aza-MLN-4760 bound to hACE2 confirmed a similar binding mode of these derivatives to the previously published binding mode of MLN-4760 [26]. The X-ray diffraction analysis of the co-crystallized hACE2/F-MLN-4760 further corroborated the analogous binding mode of F-MLN-4760 and MLN-4760 characterized by direct interactions between the carboxylate group of these compounds and the zinc ion site in a closed conformation of the hACE2 peptidase domain (Fig. 3). The X-ray diffraction analysis confirmed an equal configuration of the F-MLN-4760 as for the known (*S,S*)-MLN-4760, indicating that the (*S,S*)-diastereoisomer of the synthesized F-MLN-4760 was correctly identified.

**Fig. 3.**
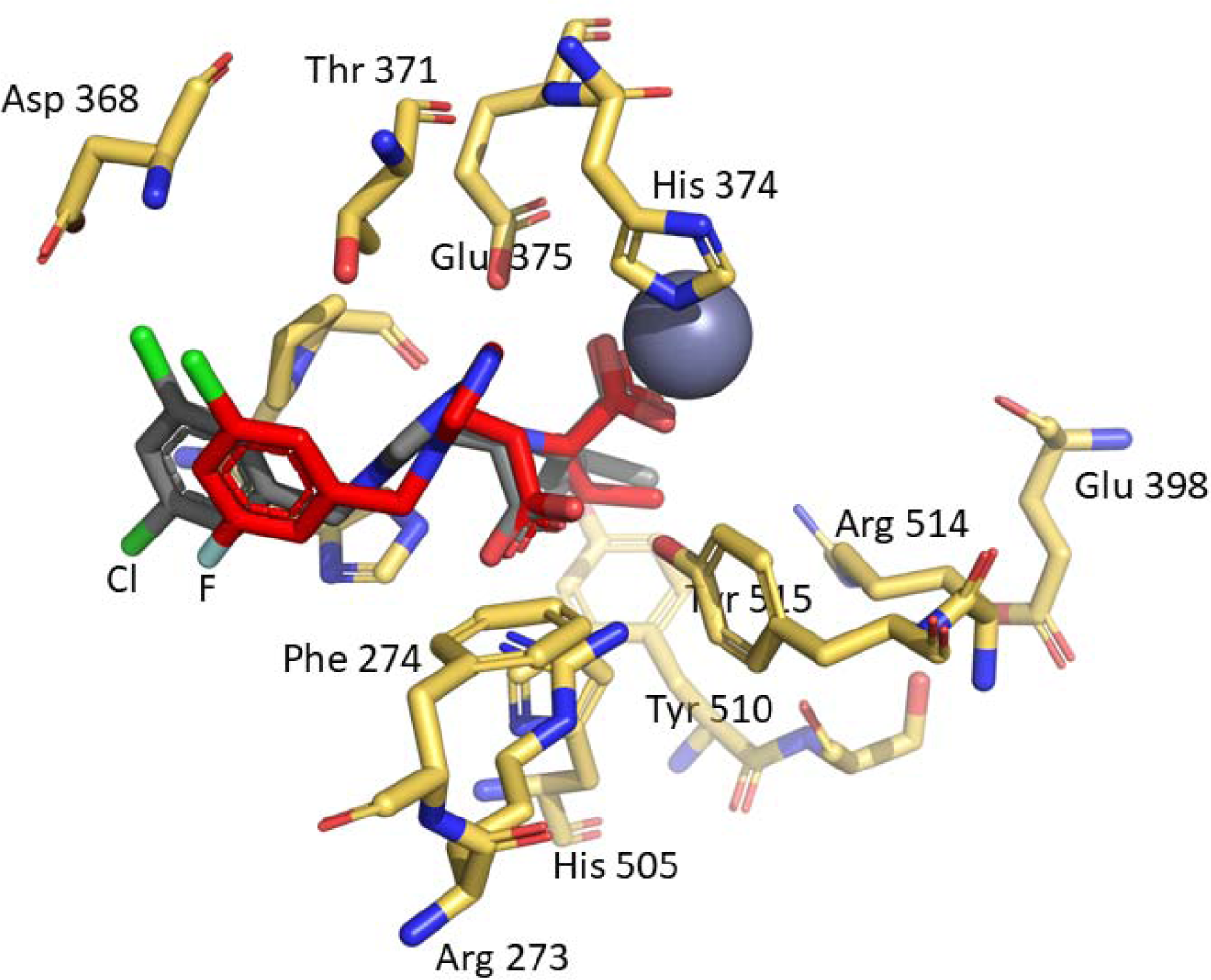
Superimposition of the hACE2/MLN-4760 complex (PDB:1R4L [26]) on the chain A of the hACE2/F-MLN-4760 (this work). A selection of key hACE2 residues at the binding site is shown with yellow carbon atoms and only for the hACE2/F-MLN-4760 complex. MLN-4760 is shown in grey and the fluorinated compound (F-MLN-4760) in red (atom colors: N = blue, O = red, Cl = light green, F = light blue). The figure was created using PyMOL (The PyMOL Molecular Graphics System, Version 2.5.3, Schrödinger LLC).

### Synthesis of MLN-4760, F-MLN-4760 and F-Aza-MLN-4760

MLN-4760, F-MLN-4760 and F-Aza-MLN-4760 were obtained in overall yields of 7%, 5% and 7%, respectively, and a purity of >99%. HRMS and NMR data correlated well with the theoretical values, confirming the structural identity of the obtained compounds (Supplementary Material).

### Synthesis of the MLN-4760-based precursors for radiofluorination

Precursors **1** and **2** were obtained with a purity of >99% and an overall yield of 8% and 28%, respectively (Supplementary Material). The carboxylic functions of both precursors were protected by *tert*-butyl esters. In precursor **1**, one of the chloride atoms of the MLN-4760 structure was replaced with a SnMe_3_-leaving group to enable radiofluorination via a copper-catalyzed reaction to obtain [^18^F]F-MLN-4760. The benzene-to-pyridine substitution in precursor **2** was suitable for radiofluorination by a halogen exchange reaction to obtain [^18^F]F-Aza-MLN-4760.

### Radiochemical synthesis, stability and n-octanol/PBS distribution coefficients

[^18^F]F-MLN-4760 and [^18^F]F-Aza-MLN-4760 were obtained with radiochemical purities of >99% and a decay-corrected radiochemical yield of 5.3% and 1.2%, respectively. The molar activities of [^18^F]F-MLN-4760 and [^18^F]F-Aza-MLN-4760 ranged from 21 to 38 GBq/µmol (n = 5) and from 78 to 81 GBq/µmol (n = 3) at the end of the synthesis, respectively. The chemical identity of the final products was confirmed by congruent UV-HPLC and radio-HPLC chromatogram peaks obtained during the co-injection of individual radiotracers with the respective (*S,S*)-diastereomers of the non-radioactive reference compound (Supplementary Material, Fig. S2 and S3). In the final formulation, both [^18^F]F-MLN-4760 and [^18^F]F-Aza-MLN-4760 were entirely stable (>97% intact radiotracer) for at least 3 h at RT. Both radiotracers were also metabolically stabe in vitro, demonstrated by >97% intact radiotracer after 3-h incubation period in murine and human blood plasma at 37°C (Supplementary Material, Fig. S4).

The logD values of the two radiotracers were relatively high with [^18^F]F-MLN-4760 (logD: −1.32 ± 0.04, n=3) being slightly more lipophilic than [^18^F]F-Aza-MLN-4760 (logD: −2.02 ± 0.21 n=3).

### ACE2-affinity of the fluorinated MLN-4760 derivatives relative to MLN-4760

The IC_50_ values of F-MLN-4760 and F-Aza-MLN-4760, obtained under the given experimental conditions, were in the high nM range similar to the value determined for MLN-4760 (Supplementary material, Fig. S5). As IC_50_ values depend on the experimental setting, the relative ACE2-binding affinities were calculated for F-MLN-4760 (0.35) and F-Aza-MLN-4760 (0.31) indicating a 3-fold and 7-fold lower affinity as compared to that of MLN-4760 (set as 1.0), for which an IC_50_ value of 0.44 nM was reported in the literature (Supplemenatery Mateiral, Fig. S5) [26].

### Uptake and internalization of the radiotracers in HEK-ACE2 and HEK-ACE cells

The HEK-ACE2 cell uptake of [^18^F]F-MLN-4760 was 49 ± 10% and 67 ± 9% after 1 h and 3 h, respectively, while that of [^18^F]F-Aza-MLN-4760 was 28 ± 10% and 37 ± 8%, respectively, after the same incubation times (Fig. 4a). Both radiotracers showed only moderate internalization (< 11%) even after a 3-h incubation period, while co-incubation of the cells with an excess of MLN-4760 blocked ACE2 and prevented the uptake of the radiotracers almost completely (<1.5%, Supplementary Material, Fig. S6). Both [^18^F]F-MLN-4760 and [^18^F]F-Aza-MLN-4760, showed only negligible uptake (<0.3%) in HEK-ACE cells even after incubation for 3 h (Fig. 4b).

**Fig. 4.**
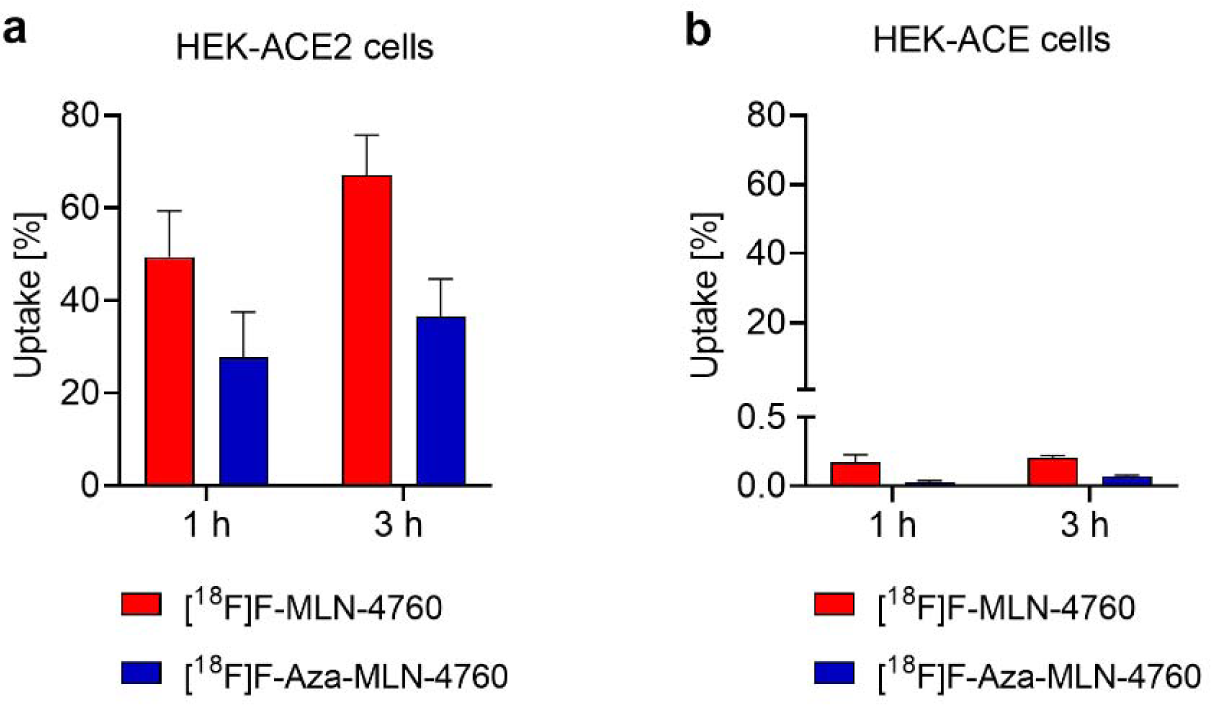
**a/b** Cell uptake of [^18^F]F-MLN-4760 and [^18^F]F-Aza-MLN-4760 after 1 h and 3 h incubation time; **a** Uptake in HEK-ACE2 cells; **b** Uptake in HEK-ACE cells.

### Autoradiography of radiotracer binding to murine tissue and HEK-xenografts

Investigation of tissue from frozen HEK-ACE2 and HEK-ACE xenografts by means of in vitro autoradiography confirmed the selectivity of [^18^F]F-MLN-4760 and [^18^F]F-Aza-MLN-4760 as negligible binding of the radiotracer to ACE-expressing tissue was observed. [^18^F]F-MLN-4760 revealed significantly higher binding to HEK-ACE2 xenografts compared to [^18^F]F-Aza-MLN-4760 (Fig. 5a). Specific binding of the [^18^F]F-MLN-4760 was observed in kidney, heart and lung tissue as well as in some parts of the brain of mice expressing murine ACE2. Quantification of the activity per area of the tissues revealed that among the physiological tissue of CD1/nude mice, the heart showed the highest radiotracer binding. Specific binding of [^18^F]F-Aza-MLN-4760 was, however, considerably lower than [^18^F]F-MLN-4760 in all represented organs (Fig. 5b). Representative autoradiograms of heart, brain, lung and kidney tissue incubated in the presence and absence of an excess of MLN-4760 can be found in the Supplementary materials (Fig S7).

**Fig. 5.**
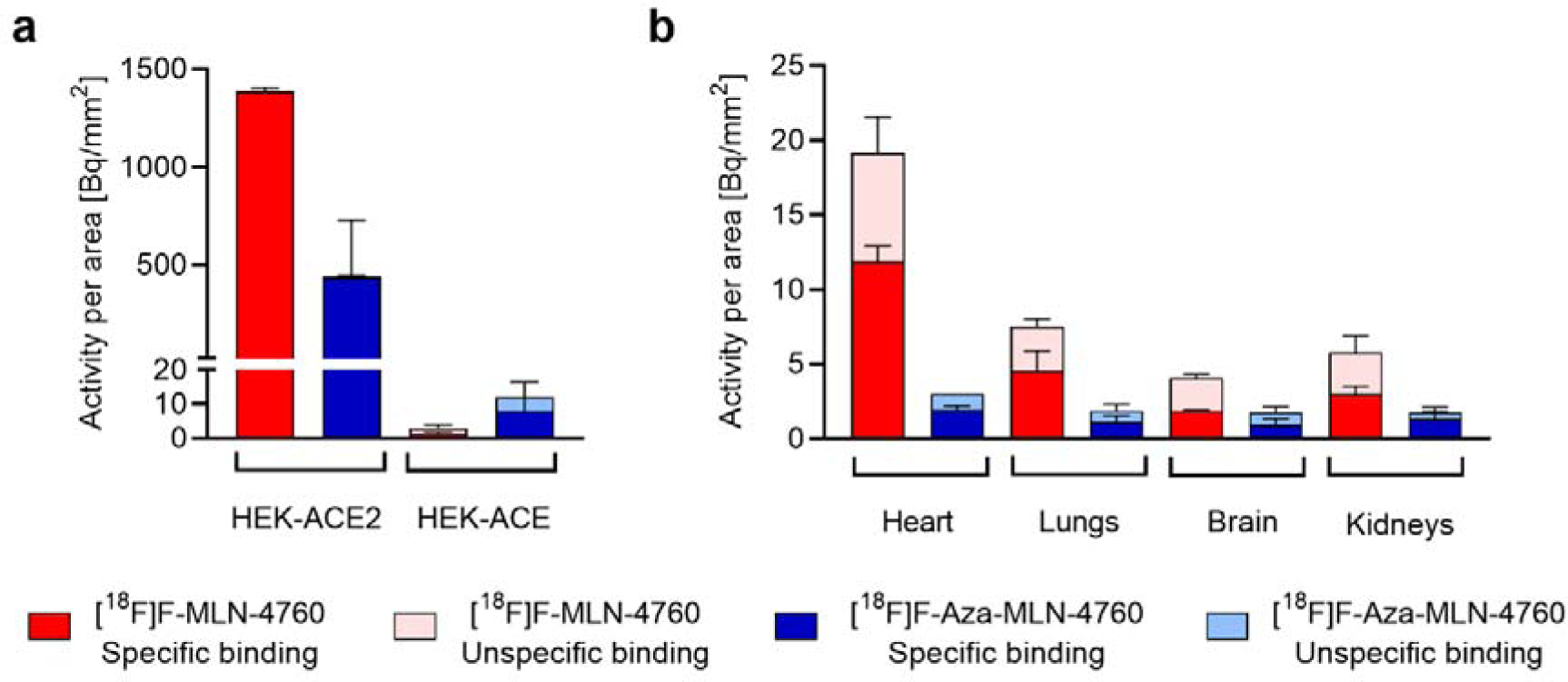
Quantification of autoradiograms performed with [^18^F]F-MLN-4760 and [^18^F]F-Aza-MLN-4760 on frozen tissue sections of mice. **a** Signal intensity obtained for HEK-ACE2 and HEK-ACE xenografts; **b** Signal intensity obtained for heart, lung, brain and kidney sections.

### Biodistribution of [^18^F]F-MLN-4760 and [^18^F]F-Aza-MLN-4760 in xenograft-bearing mice

The uptake of [^18^F]F-MLN-4760 in HEK-ACE2 xenografts reached 11 ± 1% IA/g already at 15 min p.i. and was well retained over the first hour (13 ± 2% IA/g at 1 h p.i.). At 3 h after injection of [^18^F]F-MLN-4760, still 5.8 ± 0.9% IA/g were measured in the HEK-ACE2 xenograft. A similar uptake of 15 ± 2% IA/g was obtained at 1 h after injection of [^18^F]F-Aza-MLN-4760 in the HEK-ACE2 xenografts. Activity accumulation in the HEK-ACE xenografts was negligible for both radiotracers (< 0.3% IA/g) (Fig. 6). Initial uptake of [^18^F]F-MLN-4760 in the kidneys reached 44 ± 8% IA/g (15 min p.i.), however, over the following hours, renal activity was rapidly cleared (5.1 ± 1.5% IA/g at 1 h p.i. and <0.5% IA/g at 3 h p.i.). In the case of [^18^F]F-Aza-MLN-4760, the renal uptake was 2.2 ± 0.6% IA/g at 1 h p.i. Both radiotracers showed substantial retention in the intestinal tract, reaching a maximum of 28 ± 5% IA/g and 10 ± 5% IA/g at 1 h after injection of [^18^F]F-MLN-4760 and [^18^F]F-Aza-MLN-4760, respectively ((Fig. 6; Supplementary Material, Table S1).

**Fig. 6.**
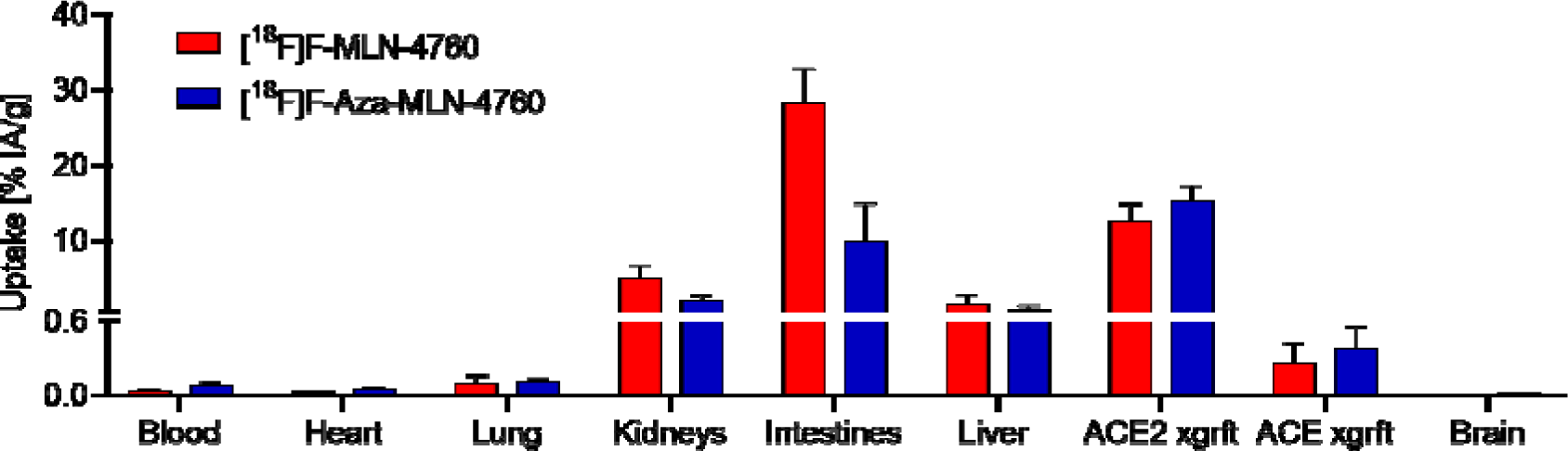
Decay-corrected biodistribution data of [^18^F]F-MLN-4760 and [^18^F]F-Aza-MLN-4760 in xenografted CD1 nude mice (n=3) 1 h after injection of the respective radiotracer.

### PET/CT studies

PET/CT imaging studies performed with mice bearing HEK-ACE2 and HEK-ACE xenografts visualized the results of the quantitative biodistribution data of [^18^F]F-MLN-4760 and [^18^F]F-Aza-MLN-4760 (Fig. 7a/b). While an intense signal was obtained from the HEK-ACE2 xenografts already at early timepoints after injection of [^18^F]F-MLN-4760 and [^18^F]F-Aza-MLN-4760, no activity accumulation was seen in the HEK-ACE xenografts. The PET images revealed substantial activity in the kidneys at early timepoints, which was entirely cleared over the first hour after injection. The highest activity retention of both radiotracers was found in the gallbladder and the intestines throughout the entire time of investigation.

**Fig. 7.**
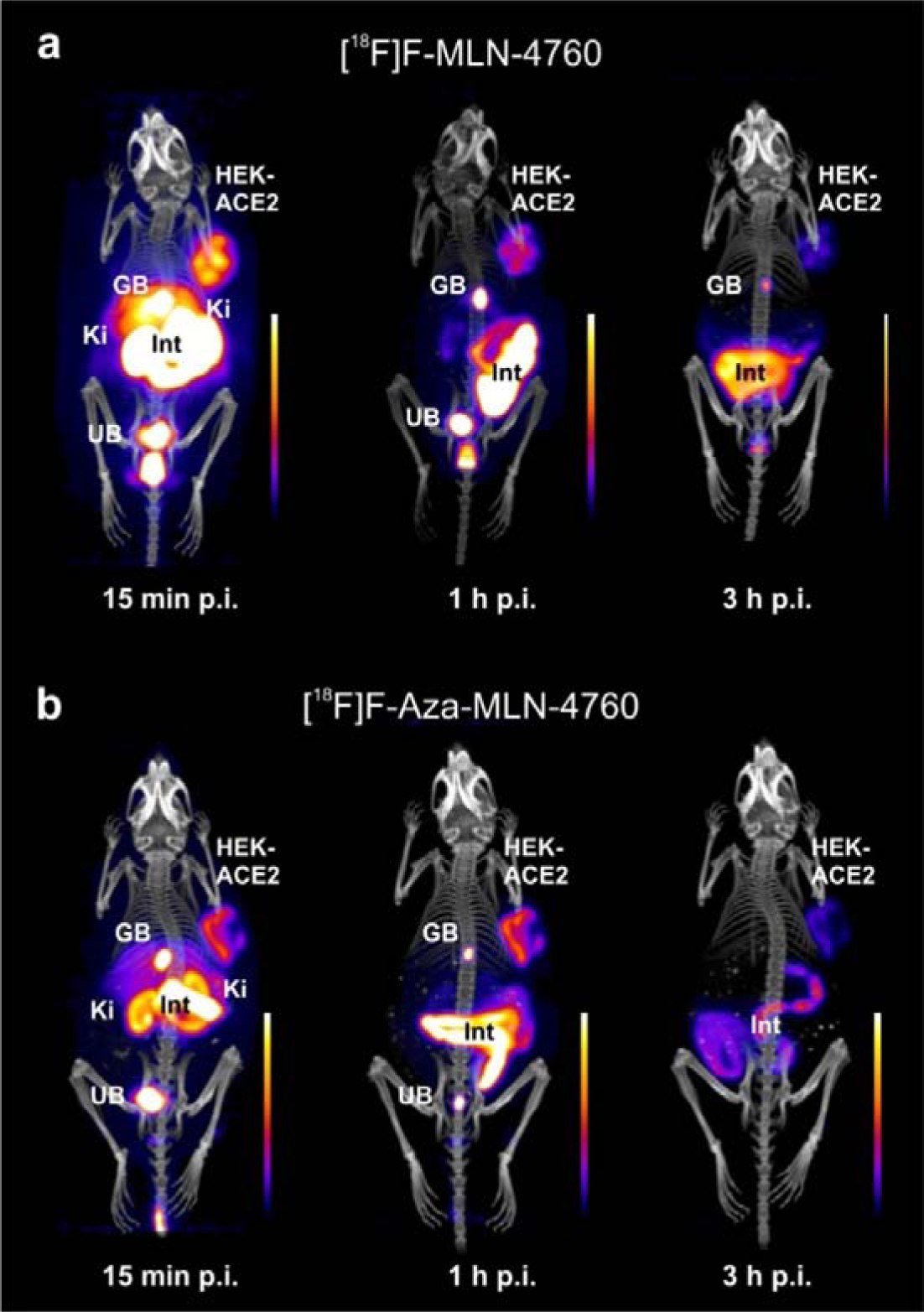
**a/b** PET/CT images of mice injected with the radiotracers. **a** PET/CT images acquired 15 min, 1 h and 3 h after injection of [^18^F]F-MLN-4760 in nude mice bearing HEK-ACE2 xenografts on the right shoulder (and HEK-ACE xenografts on the left shoulder). **b** PET/CT images acquired 15 min, 1 h and 3 h after injection of [^18^F]F-Aza-MLN-4760 in nude mice bearing HEK-ACE2 xenografts on the right shoulder (and HEK-ACE xenografts on the left shoulder). (HEK-ACE2, HEK-ACE2 xenograft; GB, gall bladder; Ki = kidney, Int, intestines; UB, urinary bladder).

## Discussion

In this study, two ^18^F-based radiotracers were developed based on the structure of MLN-4760. Computer generated models suggested that the replacement of the chlorinated benzene ring of MLN-4760 with a fluorophenyl (F-MLN-4760) or fluoropyridine (F-Aza-MLN-4760) entity would not substantially affect the ACE2-binding mode. In order to confirm the maintained ACE2-binding mode, the reference compounds were synthesized. F-MLN-4760 was prepared according to previously reported procedures [23, 33], while a new synthetic pathway involving *tert*-butyl protection of the carboxylic functions was applied for the preparation of F-Aza-MLN-4760. In both cases, the biologically active (*S,S*)-diastereoisomers were identified based on the analogous HPLC elution sequence of the diastereoismers of MLN-4760 of which the retention time of the (*S,S*)-diastereoisomer was verified using the commercially available (*S,S*)-MLN-4760. The co-crystallization of F-MLN-4760 with hACE2 confirmed the correct (*S,S*)-configuration of the stereocenters of F-MLN-4760 and its conserved binding mode to the enzyme. Moreover, the obtained data highlighted that the *para*-position of the benzene ring does not have direct interaction with hACE2 and, hence, the insertion of a nitrogen atom in this position would not affect the binding mode of the resulting pyridine-based compound, F-Aza-MLN-4760. This was further corroborated in preclinical in vitro displacement studies using [^3^H]MLN-4760, which showed that the affinity of F-MLN-4760 and F-Aza-MLN-4760 to HEK-ACE2 cells was in a similar range, albeit 3-fold and 7-fold lower than that of MLN-4760.

The preparation of the precursors of [^18^F]F-MLN-4760 and [^18^F]F-Aza-MLN-4760, respectively, was based on the synthesis pathway of F-Aza-MLN-4760 with minor modifications. The introduction of *tert*-butyl protecting groups for the carboxylic acid functionalities of the MLN-4760 scaffold allowed deprotection after radiofluorination under relatively mild conditions. As a result, the radiotracers were obtained in high yields within a short time. The design of [^18^F]F-Aza-MLN-4760 was based on the assumption that the radiofluorination of a pyridine-based precursor would enable a straightforward S_N_Ar reaction while avoiding the handling of toxic reagents required for the production of the stannylated precursor for the production of [^18^F]F-MLN-4760. Nevertheless, [^18^F]F-MLN-4760 was obtained in a 4-fold higher radiochemical yield than [^18^F]F-Aza-MLN-4760 and within a shorter period. The achieved molar activity of [^18^F]F-Aza-MLN-4760 was, however, about double as high as that of [^18^F]F-MLN-4760 which may be relevant to visualizing ACE2 at low expression levels.

Preclinical investigations confirmed the specific binding of both radiotracers to HEK-ACE2 cells in vitro and selective binding to ACE2-expressing xenografts in vivo, while almost no uptake was seen in ACE-expressing cells and xenografts. These findings are relevant as the developed PET agent should finally serve for specific and selective imaging of ACE2 without cross reacting with ACE. In vitro autoradiography studies demonstrated the feasibility of imaging ACE2 expression at physiological expression levels in murine tissue. The signals in autoradiograms were indeed congruent with the ACE2 tissue expression reported in the literature [36]. [^18^F]F-Aza-MLN-4760 showed, however, a lower signal on autoradiograms as well as lower uptake in HEK-ACE2 cells in vitro. This could be ascribed to the approximately 4-fold weaker ACE2-binding affinity of [^18^F]F-Aza-MLN-4760 as compared to the ACE2 affinity of [^18^F]F-MLN-4760. Even though the binding affinity did not seem to affect the accumulation in the xenografts in vivo, which was similar for both radiotracers, the higher binding affinity of [^18^F]F-MLN-4760 would certainly be essential in view of the imaging of considerably lower levels of ACE2 under (patho)physiological conditions.

Compared to the previously developed ACE2-targeting radiopeptides [22], a relatively high retention of activity was seen in the gall bladder and intestinal tract that can be ascribed to the rather lipophilic properties of the radiotracers, consistent with their relatively high logD. Although the initial signal in the kidneys was quite substantial, renal clearance was rather fast, resulting in almost background levels of retained activity in the kidney at 3 h p.i.. Retention in other organs and tissues was generally low.

## Conclusion

Based on various criteria, including the higher radiosynthesis yield and increased ACE2-binding affinity, [^18^F]F-MLN-4760 evolved as the preferred candidate for further preclinical investigations and potential translation to the clinics. Such a radiotracer may not only serve to image ACE2 expression dynamics during and after Covid-19, but also be a useful tool to investigate the role of ACE2 in cardiovascular diseases. A better understanding of the ACE2 regulatory function as part of the RAAS may serve future drug development processes for the treatment of related diseases.

## Supplementary Information

Available.

## Acknowlegdment

The authors thank Susan Cohrs and Fan Sozzi-Guo for technical assistance with the experiments at PSI and Annette Krämer, Henrik Mikael Portmann and Bruno Mancosu for technical assistance of the experiments at ETH Zurich. The authors thank Mara Wieser for support in protein production and crystallization at PSI, and Matthew Rodrigues for advice and support on ligand geometry refinement in the crystal structure. The authors thank Luisa M. Deberle for general support of the project at ETH.

## Funding

The entire project and in particular Darja Beyer, Jingling Wang and Yingfang He were supported by the Swiss National Science Foundation (SNSF), as a part of the National Research Program 78 (NRP78: Grant N° 4078P0_198274). Darja Beyer was additionally funded by another grant of the Swiss National Science Foundation (Grant N° 310030_188978). Christian Vaccarin was funded by the Swiss National Science Foundation (Grant N° 210456).

## Data availability

The raw data of the results presented in this study are available on request from the corresponding author.

## Declarations

### Ethics approval

This study was performed in agreement with the national law and PSI-internal guidelines of radiation safety protection. In vivo experiments were approved by the local veterinarian department and ethics committee and conducted in accordance with the Swiss law of animal protection.

### Conflict of interests

The authors have no relevant financial or non-financial interests to disclose.

## Notes

### Competing Interest Statement

The authors have declared no competing interest.

