## Supplementary Material for "Development of radiofluorinated MLN-4760 derivatives for PET imaging of the SARS-CoV-2 entry receptor ACE2"

**1. Molecular computer-based model to predict the binding mode of the MLN-4760 derivatives**

**Purpose:** A computational structural model was developed based on the published crystallographic structure of the human ACE2 (hACE2)/MLN-4760 complex [1] to predict the binding mode of the fluorinated MLN-4760 derivatives F-MLN-4760 and F-Aza-MLN-4760.

**Methods:** Using PyMOL (The PyMOL Molecular Graphics System, Version 2.5.4, Schrödinger, LLC), the published structure of the hACE2/MLN-4760 complex was modified as follows. First, the MLN-4760 was modified to build the derivatives F-MLN-4760 and F-Aza-MLN-4760. Then, the geometry of binding pose of each derivative was optimized within the binding pocket of hACE2.

**Results**: The binding modes of F-MLN-4760 and F-Aza-MLN-4760 to hACE2 were predicted to be very similar to that of MLN-4760 (Fig. S1). In both derivatives, the zinc-bound carboxylate group would mimic the tetrahedral intermediate characteristic of nucleophilic attack during peptide hydrolysis. However, the models suggested slight differences in the position of the fluorinated benzylimidazole group of both F-MLN-4760 and F-Aza-MLN-4760 with respect to the reference compound. Specifically, we observed a slight relocation of the fluorinated benzylimidazole that might result in a somewhat different set of interaction with the nearby residues Asp368, Asp367, Asn149, and His145 of hACE2.

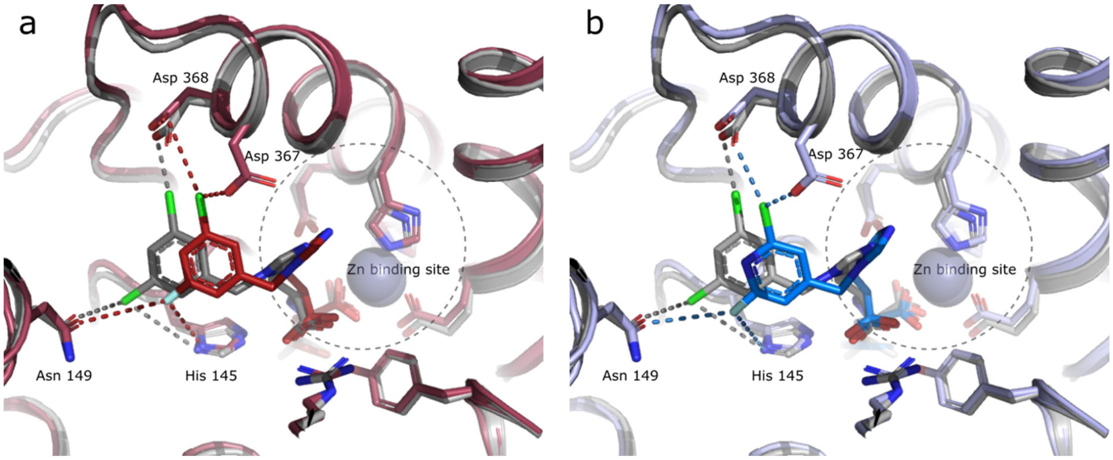


**Fig. S1 a** Comparison of the hACE2-binding modes of MLN-4760 (grey) and F-MLN-4760 (red). The protein structure is shown as ribbons and the inhibitors and selected side chains in the binding pocket are shown as sticks (atom colors: N = blue, O = red, Cl = light green, F = light blue). The distances between the halogen atoms and their interacting partners are shown as dashes. **b** Comparison of the hACE2-binding modes of MLN-4760 (grey) and F-Aza-MLN-4760 (blue).

### 2. Co-crystallization and X-ray diffraction of the F-MLN-4760/hACE2 complex

**Purpose:** The binding mode of the synthesized F-MLN-4760 to hACE2 was investigated based on the F-MLN-4760/ACE2 co-crystallization using X-ray diffraction experiments.

**Methods**: The construct comprising the extracellular part of hACE2 and a C-terminal His-tag was expressed in insect cells by secretion into the medium and purified from the medium by nickel-nitrilotriacetic acid (Ni-NTA) immobilized-metal affinity chromatography (IMAC) and size-exclusion chromatography (SECs). The protein for crystallization was in a solution of 20 mM Tris pH 7.4, 150 mM NaCl and 20 μM $ZnCl_2$ at a concentration of 5.8 mg/mL. The F-MLN-4760 was added to the protein solution at a 35-fold molar excess. Crystals formed in a sitting drop crystallization screen (JCSG-plus HT-96 screen, Molecular Dimensions), using 100 nL protein plus 100 nL reservoir solution drops. The crystallization condition was 0.1 M sodium citrate pH 5.5, 20% w/v PEG 3000. The crystals were optimized by manually setting up additional drops using the same conditions. Ethylene glycol was added as a cryoprotecting agent to a final concentration of 45%. A dataset to 2.5 Å was collected from a single crystal at 100 K at the X06SA beamline of the Swiss Light Source synchrotron (Paul Scherrer Institute in Villgen-PSI, Switzerland), using a wavelength of 1.0 Å. The data were processed using XDS software [2]. The space group of the crystal was P 1 21 1. The structure was solved by molecular replacement with Phaser [3], using an AlphaFold [4] model of hACE2 as the search model. There were two hACE2 molecules in the asymmetric unit. The initial model from molecular replacement was refined in iterative steps of model building [5] and refinement [6]. MolProbity [7] was used to validate the geometry and stereochemistry of the protein chains in the crystal structure.

**Results:** The result is reported in the main article.

### 3. Synthesis of MLN-4760, F-MLN-4760 and F-Aza-MLN-4760

#### 3.1. Materials

All commercial chemicals and solvents (Merck, TCI, Fluorochem) were used without further purification. Analytical thin-layer chromatography (TLC) was performed on pre-coated silica gel plates (Merck 60-F-254, 0.25 mm). Column chromatography was performed on silica gel (Supelco, silica gel high-purity grade (9385)), eluting the compounds with the reported solvent mixtures. Nuclear magnetic resonance (NMR) spectra were recorded on a Bruker 400 MHz (Bruker AvanceCore 400) or 500 MHz (Bruker Ascend 500) spectrometers using a solution of the tested compound in the reported deuterated solvent (Merck, *d* >99%) and using the solvents residual peaks as internal standard. Coupling constants are given in Hz and chemical shifts (δ) are reported in parts per million (ppm), relative to the residual solvent peak or tetramethylsilane (TMS). Infrared (IR) spectra were acquired in a JASCO FT-IR-4100 spectrometer and reported as wavenumber in $cm^{-1}$. Optical rotation (α) were measured on a JACSO P-2000 polarimeter. High resolution mass spectrometry (HRMS) spectra were acquired using an Acquity UPLC system equipped with a Waters Xevo Q-TOF ESI (Waters, Milford, US). Reported yields referred to the purified compounds and reaction conditions were not optimized. The purity of the synthesized compounds was determined using LCMS (Brucker Amazon1 LCMS-system connected to Agilent 1290 infinity II).

#### 3.2. Synthesis of MLN-4760 and F-MLN-4760

**Purpose:** The lead compound MLN-4760 was synthesized as a reference compound to determine its ACE2-binding affinity under the same conditions as the fluorinated derivatives. The F-MLN-4760 was synthesized as a reference compound for the radiosynthesis and for determination of its ACE2-binding affinity.

**Methods:** MLN-4760 was synthesized according to previously published procedures [8, 9]. The fluorinated derivative F-MLN-4760 was synthesized following a synthesis approach with slight modifications (Scheme S1). All synthesis intermediates and final compounds were fully characterized by $^{1}$H NMR, $^{13}$C NMR and HRMS.

**Scheme S1.** Synthesis scheme of MLN-4760 and F-MLN-4760

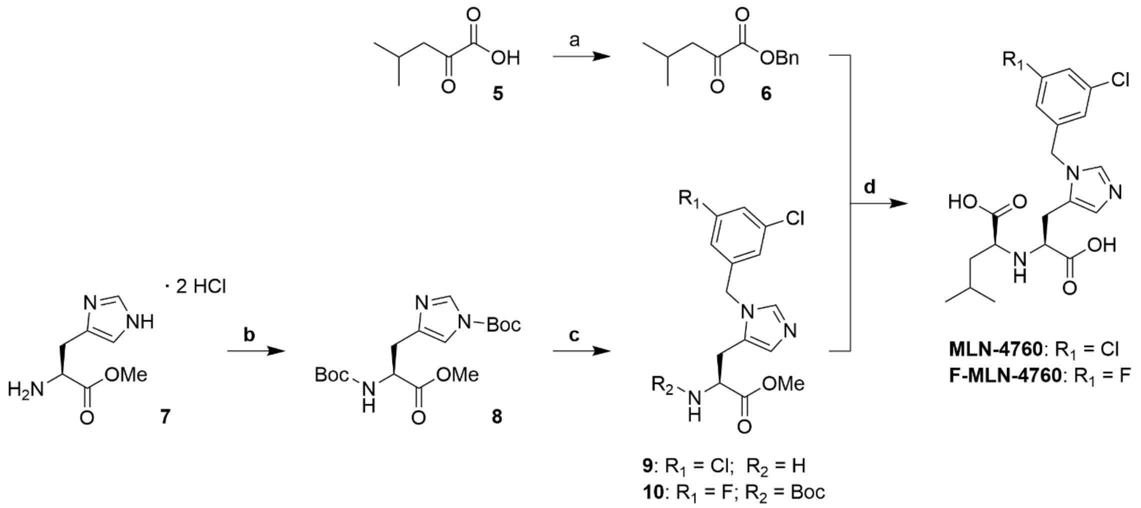


Reaction conditions: (**a**) BnOH, MsCl, pyridine, THF, 0 °C to RT, overnight; (**b**) $Boc_2O$, TEA, MeOH, 0 °C to RT, overnight; (**c**) 3,5-dicholorobenzyl alcohol or (3-chloro-5-fluorophenyl)methanol, $Tf_2O$, DIPEA, DCM, –78 °C to RT, overnight; (**d**) 1. HCl 4 M in dioxane, RT, 2 h; 2. $NaB(OAc)_3H$, DCE, RT, overnight; 3. $NaHCO_3$, RT, 1 h; 4. NaOH 1 M (aq), MeOH, RT, 1 h.

***Synthesis of intermediate 6***. 4-Methyl-2-oxopentanic acid (**5**, 950 µL, 7.68 mmol, 1.00 equiv), benzyl alcohol (1.59 mL, 15.4 mmol, 2.00 equiv) and pyridine (1.55 mL, 19.2 mmol. 2.50 equiv) were dissolved in tetrahydrofurane (THF, 8 mL) and cooled to 0 °C. Following the dropwise addition of methanesulfyonyl chloride (720 µL, 9.22 mmol. 1.20 equiv), the reaction mixture was stirred at room temperature (RT) overnight. The reaction mixture was quenched with deionized $H_2O$ (16 mL) and extracted with diethyl ether ($Et_2O$, 3 x 16 mL). The combined organic layers were dried over $Na_2SO_4$ and concentrated under reduced pressure. The resulting oil was purified by column chromatography (10:1 to 1:1 hexane:ethyl acetate (EtOAc)), to obtain intermediate **6** (1.56 g, 92% yield).

***Synthesis of intermediate 8.*** Di-*tert*-butyl dicarbonate (11.9 mL, 51.6 mmol, 2.50 equiv) was dissolved in 6 mL methanol (MeOH). The mixture was slowly added to a cooled (0 °C) solution of (*S*)-histidine methyl ester (5.00 g, 20.6 mmol, 1.00 equiv) and triethylamine (11.5 mL, 82.6 mmol, 4.00 equiv) in MeOH (52 mL). The reaction was stirred at RT overnight, after which it was concentrated under reduced pressure. The resulting residue was redissolved in dichloromethane (DCM, 20 mL), washed with deionized $H_2O$ (3 x 20 mL), dried over $Na_2SO_4$ and concentrated under vacuum. The resulting oil was triturated with cold hexane to provide compound **8** as a white solid (4.71 g, 62% yield).

***Synthesis of intermediate 9***. *N,N*-diisopropylethylamine (DIPEA, 1.88 mL, 10.7 mmol 1.10 equiv) was added to a solution of 3,5-dichlorobenzyl alcohol (1.90 g, 10.7 mmol, 1.10 equiv) in DCM (9 mL) followed by cooling to –78 °C under argon atmosphere. A solution of trifluoromethanesulfonic anhydride (1.81 mL, 10.7 mmol, 1.10 equiv) in DCM (36 mL) was added under argon and the resulting mixture was stirred for 30 min. Subsequently, a solution of compound **8** (3.60 g, 9.76 mmol, 1.00 equiv) in DCM (10 mL) was added and the mixture was stirred at RT overnight. The resulting mixture was washed with saturated aq. $NaHCO_3$ (3 x 15 mL), followed by brine (1 x 15 mL) and dried over $Na_2SO_4$. The crude product was purified by column chromatography (1-5% MeOH in DCM). The Boc group was deprotected by stirring the compound in 4 M HCl in dioxane (13.4 mL, 53.7 mmol, 23.0 equiv.) at RT for 1.5 h. Subsequently, the solvent was evaporated and the resulting di-hydrochloride salt suspended in EtOAc (10 mL). The suspension was filtered and the recovered solid was dried under high vacuum for 2 h to afford intermediate **9** as a white dihydrochloride salt (960 mg, 30% yield.)

***Synthesis of intermediate 10***. DIPEA (1.13 mL, 6.25 mmol, 1.10 equiv) was added to a solution of (3-chloro-5-fluorophenyl)methanol (1.00 g, 6.25 mmol, 1.10 equiv) in DCM (5 mL) followed by cooling to –78 °C. A solution of trifluoromethanesulfonic anhydride (1.06 mL, 6.25 mmol, 1.10 equiv) in DCM (21 mL) was added under argon and the resulting mixture was stirred at –78 °C for 30 min. Subsequently, a solution of compound **8** (2.10 g, 5.68 mmol, 1.00 equiv) in DCM (6 mL) was added, and the mixture was stirred at RT overnight. The crude reaction mixture was washed with aq. $NaHCO_3$ (3 x 15 mL), brine (1 x 15 mL), and dried over $Na_2SO_4$. The crude product was purified using silica gel chromatography (1-5% MeOH in DCM) to afford compound **10** as a yellow oil (787 mg, 34% yield).

***Synthesis of MLN-4760***. Intermediate **9** (549 mg, 1.15 mmol, 1.00 equiv) was suspended in 1,2-dichloroethane (11 mL), the $\alpha$-ketoester **6** (504 mg, 2.30 mmol, 2.00 equiv) was added to the suspension and the reaction mixture was left to stir for 2.5 h. $NaB(OAc)_3H$ (728 mg, 3.45 mmol, 3.00 equiv) was added slowly and the resulting solution was stirred overnight. The pH of the solution was adjusted to pH 8 with saturated aq. $NaHCO_3$ and the mixture was stirred for 1 h. The phases were separated and the aqueous layer was extracted with EtOAc (2 x 30 mL). The combined organic layers were dried over

$Na_2SO_4$, filtered, and concentrated to give a yellow oil. The crude product was purified by column chromatography (hexanes:EtOAc 4:1 (*v/v*), then EtOAc, then 5% MeOH in EtOAc) to afford the diastereomeric mixture as a yellow oil. A solution of 1 M NaOH (6.90 mL, 6.90 mmol, 6.00 equiv.) and MeOH (6 mL) was added to the crude product and stirred for 1 h. The deprotected diastereomers were separated using RP-HPLC (5–75% acetonitrile (MeCN) in MilliQ $H_2O$ with 0.1% TFA). The (*S,S*)-diasteromer of MLN-4760 was identified based on the HPLC elution profile of the commercially available enantiopure (*S,S*)-MLN-4760 (MW: 428.31; Merck, CAS N° 305335-31-3) as a reference. The collected fractions were then lyophilized to afford the pure (*S,S*)-diastereomer of **MLN-4760** as a white solid (215 mg, 43% yield).

***Synthesis of F-MLN-4760***. Intermediate **10** (495 mg, 1.20 mmol, 1.00 equiv) was treated with 4 M HCl in dioxane (6.92 mL, 23.0 equiv) for 1.5 h. The solvent was evaporated and the resultant compound, obtained as a di-hydrochloride salt, was suspended in EtOAc (10 mL). The suspension was filtered and the recovered solid was dried under high vacuum for 2 h. The remaining off-white solid (332 mg, 0.87 mmol, 1.00 equiv) was suspended in 1,2-dichloroethane (9 mL), the ketoester **6** (383 mg, 1.74 mmol, 2.00 equiv) was added to the suspension and the reaction mixture was left to stir for 2.5 h. $NaB(OAc)_3H$ (553 mg, 2.61 mmol, 3.00 equiv) was added slowly and the resulting solution was stirred overnight. The pH of the solution was adjusted to pH 8 with saturated aq. $NaHCO_3$ and the mixture was stirred for 1 h. The phases were separated and the aqueous layer was extracted with EtOAc (2 x 30 mL). The combined organic layers were dried over $Na_2SO_4$, filtered and concentrated to give a yellow oil. The crude product was purified by silica gel chromatography (hexanes:EtOAc 4:1 (*v/v*), then EtOAc, then 5% MeOH in EtOAc) to obtain the diastereomeric mixture as a yellow oil. Subsequently, a solution of 1 M NaOH (5.22 mL, 5.22 mmol, 6.00 equiv) and MeOH (5 mL) was added to the crude product and stirred for 1 h. The deprotected diastereomers were separated with RP-HPLC (5-75% MeCN in MilliQ $H_2O$ + 0.1% TFA). The desired (*S,S*)-diasteromer was identified based on the assumption that the HPLC elution profile would be similar to that of MLN-4760 of which the enantiopure (*S,S*)-MLN-4760 was commercially available. The desired (*S,S*)-diastereomer of F-MLN-4760 was identified based on the assumption that the sequence of the eluted diastereoisomers would be the same as that of the diastereoisomers of MLN-4760. The collected fractions were then lyophilized to afford the pure (*S,S*)-diastereomer of **F-MLN-4760** as a white solid (124 mg, 28% yield).

**Results:** The synthesis intermediates and final compounds were chemically characterized by $^1$H and $^{13}$C NMR spectra and HRMS data and for some compounds, also IR data were acquired.

***Characterization of intermediate 6***: **$^1$H NMR** (500 MHz, $CD_2Cl_2$) δ[ppm] 7.52 – 7.31 (m, 5H), 5.32 (d, $J$ = 0.9 Hz, 2H), 2.78 (dd, $J$ = 6.8, 0.8 Hz, 2H), 2.28 – 2.16 (m, 1H), 1.01 (dt, $J$ = 6.7, 0.8 Hz, 6H). **$^{13}$C NMR** (126 MHz, $CD_2Cl_2$) δ[ppm] 194.33, 161.46, 135.25, 129.10, 129.08, 128.94, 68.15, 48.34, 24.48, 22.55. **HRMS** (ESI): calculated for $C_{13}H_{16}O_3$ $[M+Na]^+$:243.0992, found: 243.0992.

***Characterization of intermediate 8***: $^{1}$**H NMR** (400 MHz, $CD_2Cl_2$) δ[ppm] 7.94 (d, *J* = 1.4 Hz, 1H), 7.13 (d, *J* = 1.3 Hz, 1H), 5.92 (d, *J* = 8.4 Hz, 1H), 4.48 (dt, *J* = 8.5, 5.5 Hz, 1H), 3.65 (s, 3H), 3.00 – 2.91 (m, 2H), 1.53 (s, 9H), 1.37 (s, 9H). $^{13}$**C NMR** (126 MHz, $CD_2Cl_2$) δ[ppm] 172.81, 155.89, 147.52, 139.45, 137.47, 115.14, 85.99, 79.86, 55.23, 54.48, 53.84, 52.60, 30.68, 28.66, 28.18. **HRMS** (ESI): calculated for $C_{17}H_{27}N_3O_6$ $[M+Na]^+$: 392.1792, found: 392.1797.

***Characterization of intermediate 9***: $^{1}$**H NMR** (500 MHz, $D_2O$) δ[ppm] 8.93 (d, *J* = 1.5 Hz, 1H), 7.59 (d, *J* = 1.4 Hz, 1H), 7.58 – 7.54 (m, 1H), 7.30 (dd, *J* = 2.0, 1.0 Hz, 2H), 5.51 (s, 2H), 4.17 (td, *J* = 7.3, 1.3 Hz, 1H), 3.82 (d, *J* = 0.8 Hz, 3H), 3.45 – 3.24 (m, 2H). $^{13}$**C NMR** (126 MHz, $D_2O$) δ[ppm] 168.51, 136.49, 136.17, 135.53, 129.17, 128.01, 126.20 (d, *J* = 1.5 Hz), 119.88, 54.00, 50.78, 49.41, 24.00. **HRMS** (ESI): calculated for $C_{14}H_{15}Cl_2N_3O_2$ $[M+H]^+$: 328.0614, found: 328.0614.

***Characterization of intermediate 10***: $^{1}$**H NMR** (500 MHz, $CD_3OD$) δ[ppm] 8.95 (s, 1H), 7.40 (s, 1H), 7.34 – 7.18 (m, 2H), 7.18 – 6.95 (m, 1H), 5.50 (d, *J* = 1.8 Hz, 2H), 4.33 (dd, *J* = 9.2, 4.9 Hz, 1H), 3.72 (s, 3H), 3.22 – 2.86 (m, 2H), 1.37 (s, 9H). $^{13}$**C NMR** (126 MHz, $CD_3OD$) δ[ppm] 170.96, 164.14, 162.15, 161.02 (d, *J* = 35.3 Hz), 156.29, 137.54 (d, *J* = 8.5 Hz), 135.93 (d, *J* = 18.0 Hz), 131.32, 123.91, 118.83, 116.47 (d, *J* = 25.1 Hz), 113.45 (d, *J* = 23.0 Hz), 79.66, 51.77, 49.02 (d, *J* = 1.9 Hz), 27.18, 25.77. $^{19}$**F NMR** (471 MHz, $CD_3OD$) δ[ppm] -110.87. **IR** (v/cm$^{-1}$, neat): 3118.33, 2979, 2359.48, 2335.37, 1739.48, 1670.53, 1606.41, 1580.38, 1482.51, 1430.44, 1367.77, 1310.88, 1200.95, 1127.19, 1053.43. **HRMS** (ESI): calculated for $C_{19}H_{23}ClFN_3O_4$ $[M+H]^+$: 412.1434, found: 412.1425.

***Characterization of the (S,S)-diastereomer of MLN-4760***: $^{1}$**H NMR** (500 MHz, $CD_3OD$) δ[ppm] 8.99 (q, *J* = 1.8 Hz, 1H), 7.67 (d, *J* = 1.6 Hz, 1H), 7.51 (t, *J* = 1.9 Hz, 1H), 7.35 (d, *J* = 1.8 Hz, 2H), 5.53 (d, *J* = 2.6 Hz, 2H), 4.11 – 3.87 (m, 2H), 3.34 – 3.27 (m, 1H), 3.26 – 3.15 (m, 1H), 1.94 (p, *J* = 6.8 Hz, 1H), 1.83 – 1.65 (m, 2H), 0.99 (d, *J* = 6.5 Hz, 6H). $^{13}$**C NMR** (126 MHz, $CD_3OD$) δ[ppm] 173.63, 171.59, 162.80, 162.52, 138.56, 137.45, 131.36, 130.23, 127.72, 121.00, 60.42, 59.89, 50.36, 41.51, 26.53, 25.86, 22.78, 22.37. **HRMS** (ESI): calculated for $C_{19}H_{23}Cl_2N_3O_4$ $[M+H]^+$: 426.0982, found: 426.0978.

***Characterization of the (S,S)-F-MLN-4760***: $^{1}$**H NMR** (500 MHz, $CD_3OD$) δ[ppm] 9.00 (d, *J* = 1.6 Hz, 1H), 7.67 (d, *J* = 1.4 Hz, 1H), 7.29 (dt, *J* = 8.5, 2.1 Hz, 1H), 7.24 (d, *J* = 1.8 Hz, 1H), 7.09 (dt, *J* = 9.1, 1.9 Hz, 1H), 5.54 (d, *J* = 3.0 Hz, 2H), 4.09 (t, *J* = 6.8 Hz, 1H), 3.94 (dd, *J* = 7.6, 5.9 Hz, 1H), 3.30 – 3.27 (m, 1H), 3.22 (ddd, *J* = 16.1, 6.9, 1.0 Hz, 1H), 1.99 – 1.87 (m, 1H), 1.82 – 1.66 (m, 2H), 1.00 (d, *J* = 6.5 Hz, 6H). $^{13}$**C NMR** (126 MHz, $CD_3OD$) δ[ppm] 173.67, 171.52, 165.54, 163.55, 138.87, 137.49, 131.37, 125.21, 121.00, 117.97, 114.88, 60.36, 59.74, 50.47, 41.54, 26.52, 25.87, 22.77, 22.35. $^{19}$**F NMR** (471 MHz, $CD_3OD$) δ[ppm] -110.79. **IR** (v/cm$^{-1}$, neat): 3131.35, 3045.05, 2964.05, 2877.75, 2359.48, 2337.78, 1733.21, 1653.66, 1609.79, 1591.95, 1429.96, 1272.79, 1180.7, 1134.9. **HRMS** (ESI): calculated for $C_{19}H_{23}ClFN_3O_4$ $[M+H]^+$: 412.1434, found: 412.1433.

#### 3.3. Synthesis of F-Aza-MLN-4760

**Purpose:** The reference compound F-Aza-MLN-4760 was synthesized as a reference compound for the radiosynthesis and for determination of its ACE2-binding affinity.

**Methods:** F-Aza-MLN-4760 was synthesized according to the procedure reported for MLN-4760 (Section 2.1) with slight modifications (Scheme S2). The histidine alkylation was performed after the reductive amination reaction to increase the synthetic yield. All synthesis intermediates and final compounds were fully characterized by $^{1}$H NMR, $^{13}$C NMR and HRMS.

**Scheme S2.** Synthesis of F-Aza-MLN-4760

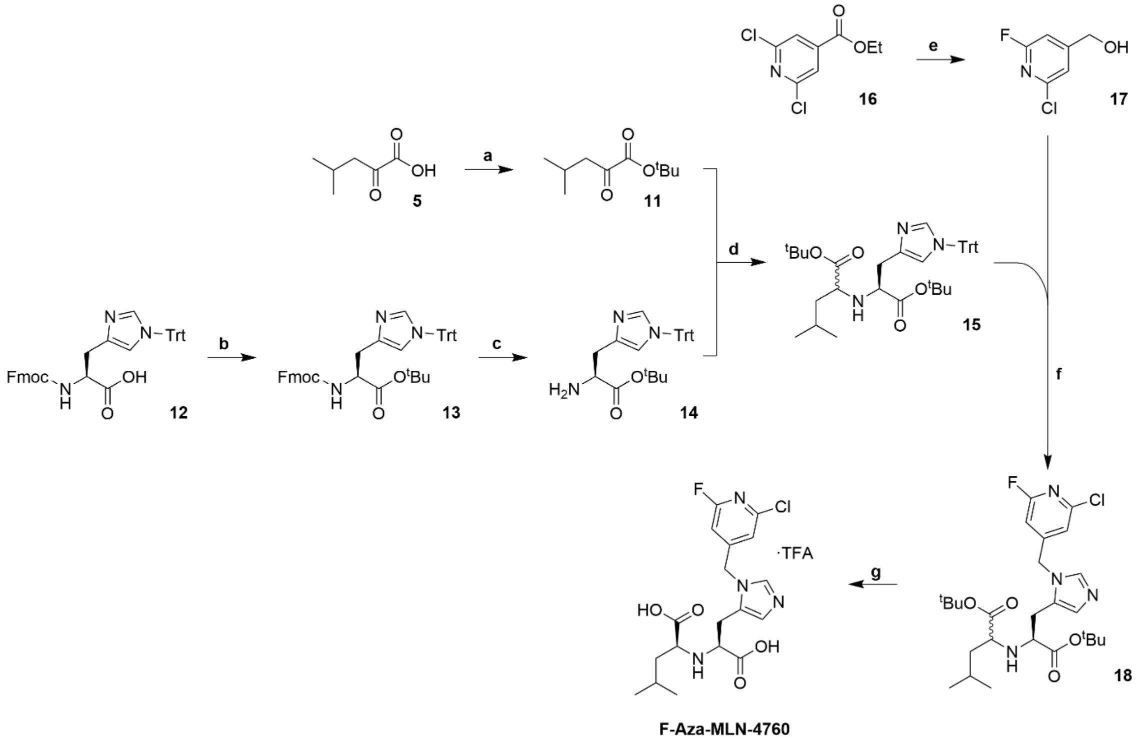


Reaction conditions: (**a**) $^{t}$BuOH, pyridine, MsCl, THF, 0 °C to RT, overnight; (**b**) *tert*-butyl-2,2,2-trichloroacetimidate, DCM, reflux, 5 h; **c**) 20% piperidine in MeCN, RT, 30 min; (**d**) 1. NaB(OAc)$_3$H, DCE, RT, overnight; 2. NaHCO$_3$, RT, 1 h; (**e**)1. CsF, DMSO, 140 °C, 1 h, 2. NaBH$_4$, MeOH, THF, 50 °C, overnight; (**f**) Tf$_2$O, DIPEA, DCM, –78 °C to RT, overnight; (**g**) TFA:MilliQ H$_2$O:TIPS (95.0:2.5:2.5, *v/v/v*), RT, 2 h.

***Synthesis of intermediate*** **11**. 4-Methyl-2-oxopentanoic acid (**5**, 950 µL, 7.70 mmol, 1.00 equiv), *tert*-butyl alcohol ($^{t}$BuOH, 1.46 mL, 15.4 mmol, 2.00 equiv) and pyridine (1.56 mL, 19.3 mmol, 2.50 equiv) were dissolved in THF (8 mL) and the reaction mixture was cooled to 0 °C. Mesyl chloride (715 µL, 9.24 mmol, 1.20 equiv) was added dropwise followed by stirring at RT overnight. The reaction was then quenched with deionized H$_2$O (16 mL) and extracted with Et$_2$O (3 x 10 mL). The combined organic layers were dried over $Na_2SO_4$, filtered, and concentrated to obtain a yellow oil. The obtained residue was purified using silica gel chromatography (hexane:EtOAc 10:1 to 2:1 (*v/v*)) to afford intermediate **11** as a yellow oil (1.10 g, 77% yield).

***Synthesis of intermediate* 13.** Fmoc-His(Trt)-OH (**12**, 5.00 g, 8.06 mmol, 1.00 equiv) was suspended in dry DCM (81 mL) and *tert*-butyl-2,2,2-trichloroacetimidate (TBTA, 2.85 mL, 16.1 mmol, 2.00 equiv) was added dropwise. The resulting mixture was refluxed for 5 h. The reaction mixture was washed with deionized $H_2O$ (3 x 80 mL) and the organic layers were dried over $Na_2SO_4$, filtered, and concentrated under reduced pressure to give a yellow oil. The crude product was purified using silica gel chromatography (hexane:EtOAc 20:1 to 1:1 (*v/v*)) to afford compound **13** as a pale yellow oil (3.00 g, 55% yield).

***Synthesis of intermediate* 14**. Compound **13** was treated with 20% piperidine in MeCN (40 mL) at RT for 30 min. Subsequently, the solvent was removed under reduced pressure and the crude product was purified with a silica gel chromatography (hexane:EtOAc 4:1 (*v/v*)), then EtOAc, then 5% MeOH in EtOAc) to obtain compound **14** as a colorless oil (1.38 g, 68% yield).

***Synthesis of intermediate* 15**. Compound **14** (210 mg, 460 μmol, 1.00 equiv) was dissolved in 1,2-dichloroethane (5 mL) and the ketoester **11** (173 mg, 930 μmol, 2.00 equiv) was added dropwise to the solution. The reaction mixture was stirred at RT for 2.5 h. $NaB(OAc)_3H$ (293 mg, 1.38 mmol, 3.00 equiv) was added slowly and the reaction was stirred at RT overnight. The pH of the solution was adjusted with aq. saturated $NaHCO_3$ to pH 8 and the resulting mixture was further stirred at RT for 1 h. The phases were separated and the aqueous layer was extracted with EtOAc (2 x 10 mL). The combined organic layers were dried over $Na_2SO_4$, filtered, and concentrated under reduced pressure. The obtained crude product was purified by silica gel chromatography (hexane/EtOAc 20:1 to 1:1 (*v/v*)) to afford a diastereomeric mixture of intermediate **15** as a yellow oil (221 mg, 77% yield, (*S,S*):(*S,R*) 3:1).

***Synthesis of intermediate* 17.** 2,6-dichloroisonicotinic acid ethyl ester (**16**, 440 mg, 2.00 mmol, 1.00 equiv) and CsF (364 mg, 2.40 mmol, 1.20 equiv) were dissolved in anhydrous DMSO (2 mL) before heating at 140 °C for 1 h under argon atmosphere. This mixture was diluted with deionized $H_2O$ (50 mL) and extracted with DCM (3 x 20 mL). The combined organic layers were dried over $Na_2SO_4$, filtered, and concentrated under reduced pressure. The obtained solid residue was dissolved in a 1:1 mixture of MeOH and THF (6 mL) and cooled to 0 °C. After the addition of $NaBH_4$ (96 mg, 3.00 mmol, 1.50 equiv) the reaction mixture was stirred at 50 °C overnight under argon atmosphere. At reaction completion, the volatile solvents were removed by evaporation under reduced pressure and the obtained residue was redissolved in 1M HCl (20 mL) before further stirring at RT for 1 h. The pH was adjusted to a value of ~8 by the addition of solid $Na_2CO_3$ and the aqueous phase was extracted with $Et_2O$ (3 x 10 mL). The combined organic layers were dried over $Na_2SO_4$, filtered, and concentrated under reduced pressure. The resulting solid was purified by RP-HPLC (15-75% MeCN in MilliQ $H_2O$ + 0.1% TFA) and the fractions containing the product of interest were collected, combined, and neutralized with aq. saturated $NaHCO_3$ to pH ~7 before proceeding with the extraction of the product with DCM (2 x 10 mL). The combined organic layers were then dried over $Na_2SO_4$ and concentrated to afford the radiofluorination precursor **17** as a white solid (78 mg, 24% yield).

***Synthesis of intermediate* 18.** DIPEA (20.9 µL, 120 µmol, 1.20 equiv) was added to a solution of compound **17** (19.2 mg, 120 µmol, 1.20 equiv) in DCM (500 µL) before cooling to –78 °C. A solution of trifluoromethanesulfonic anhydride (20.2 µL, 120 µmol, 1.20 equiv) in DCM (500 µL) was added under argon and the resulting mixture was stirred for 30 min at –78 °C followed by slow addition of compound **15** (62.0 mg, 100 µmol, 1.00 equiv) in DCM (500 µL). The reaction mixture was stirred at RT overnight. At reaction completion, the reaction mixture was diluted with DCM (10 mL), washed with deionized $H_2O$ (2 x 5 mL) and brine (5 mL), dried over $Na_2SO_4$ and concentrated by evaporation under reduced pressure. The crude was purified by silica gel chromatography (100% EtOAc, then 5% MeOH in EtOAc) to afford a diastereomeric mixture of intermediate **18** (38.0 mg, 72% yield, (*S,S*):(*S,R*) 3:1)

***Synthesis of (S,S)-F-Aza-MLN-4760.*** Compound **18** (52.0 mg, 100 µmol, 1.00 equiv) was dissolved in a mixture of TFA:MilliQ $H_2O$:triisopropylsilane (2 mL, 95.0:2.5:2.5, *v/v/v*) and then stirred at RT for 2 h. The volatile solvents were then removed by an $N_2$ stream. The deprotected diastereomers were separated with RP-HPLC (10-30% MeCN in MilliQ $H_2O$ + 0.1% TFA). The desired (*S,S*)- diastereomer of F-Aza-MLN-4760 was identified based on the assumption that the sequence of the eluted diastereoisomers would be the same as that of the diastereoisomers of MLN-4760. Fractions containing the product were lyophilized to afford the pure (*S,S*)-diastereomer of **F-Aza-MLN-4760** · TFA salt as a white solid (18.9 mg, 36% yield).

**Results:** The synthesis intermediates and final compounds were chemically characterized by $^1H$ and $^{13}C$ NMR spectra and HRMS data and for some compounds also the IR data was obtained.

***Characterization of compound 11:*** **$^1$H NMR** (500 MHz, $CDCl_3$) δ[ppm] 2.66 (ddd, *J* = 6.7, 1.3, 0.6 Hz, 2H), 2.18 (dpd, *J* = 13.4, 6.8, 1.3 Hz, 1H), 1.56 (dd, *J* = 1.3, 0.6 Hz, 9H), 0.98 (ddd, *J* = 6.7, 1.3, 0.6 Hz, 6H). **$^{13}$C NMR** (126 MHz, $CDCl_3$) δ[ppm] 195.64, 161.16, 83.94, 47.88, 27.94, 24.40, 22.62. **IR** (v/cm$^{-1}$, neat): 2961.16, 2936.09, 2872.45, 1800.22, 1717.78, 1658.48, 1574.59, 1467.08, 1394.76, 1369.21, 1313.77, 1294, 1256.88, 1164.31, 1134.9, 1050.53, 1037.52. **HRMS** (ESI): calculated for $C_{10}H_{18}O_3$ $[M+Na]^+$: 209.1148, found: 209.1152.

***Characterization of compound 13:*** **$^1$H NMR** (500 MHz, $CD_2Cl_2$) δ[ppm] 7.78 (dd, J = 7.7, 1.1 Hz, 2H), 7.64 (ddt, J = 6.8, 3.1, 1.0 Hz, 2H), 7.42 – 7.36 (m, 3H), 7.33 (dd, J = 4.9, 1.9 Hz, 8H), 7.32 – 7.26 (m, 3H), 7.16 – 7.10 (m, 6H), 6.63 (d, J = 7.7 Hz, 2H), 4.41 (dt, J = 8.3, 5.0 Hz, 1H), 4.36 – 4.20 (m, 3H), 2.99 (d, J = 5.1 Hz, 2H), 1.36 (s, 9H). **$^{13}$C NMR** (126 MHz, $CD_2Cl_2$) δ[ppm] 171.00, 156.41, 144.62, 144.53, 142.87, 141.62, 139.02, 137.00, 130.16, 128.42, 128.38, 127.98, 127.44, 127.42, 125.68, 125.65, 120.26, 119.70, 81.65, 75.60, 67.21, 54.89, 47.65, 30.30, 28.21. **IR** (v/cm-1, neat): 3323.71, 3090.85, 3059.99, 3032.51, 2976.59, 2930.79, 2898, 2361.89, 2340.19, 2248.59, 1716.34,

1509.03, 1493.6, 1477.21, 1444.9, 1392.84, 1366.8, 1323.89, 1235.18, 1218.31, 1150.33, 1080.42, 1049.09. **HRMS** (ESI): calculated for $C_{44}H_{41}N_3O_4$ $[M+H]^+$: 676.3170, found: 676.3158.

***Characterization of compound 14:*** **$^{1}$H NMR** (500 MHz, $CDCl_3$) δ[ppm] 7.34 (d, *J* = 1.4 Hz, 1H), 7.30 – 7.26 (m, 9H), 7.12 – 7.08 (m, 6H), 6.61 (dt, *J* = 1.5, 0.7 Hz, 1H), 3.66 (dd, *J* = 7.5, 4.7 Hz, 1H), 3.01 – 2.71 (m, 2H), 1.37 (s, 9H). **$^{13}$C NMR** (126 MHz, $CDCl_3$) δ[ppm] 174.19, 142.47, 138.68, 137.46, 129.75, 128.02, 119.29, 80.76, 75.15, 55.04, 33.56, 28.07. **IR** (v/cm$^{-1}$, neat): 3361.8, 2978.52, 2931.27, 2357.07, 1722.12, 1597.73, 1554.83, 1490.7, 1478.17, 1442.98, 1428.03, 1389.94, 1363.91, 1325.82, 1307.98, 1275.2, 1325.82, 1307.98, 1275.2, 1238.56, 1228.92, 1201.43, 1184.56, 1147.44, 1120.44, 1087.66, 1035.59, 1012.93, 1000.87. **HRMS** (ESI): calculated for $C_{29}H_{31}N_3O_2$ $[M+H]^+$: 454.2489, found: 454.2479.

***Characterization of compound 15:*** Compound **15** was obtained as a mixture of two diastereomers after column chromatography with a diastereomers mixture of (*S,S*):(*S,R*) 3:1 as judged per NMR. **$^{1}$H NMR** (500 MHz, $CD_2Cl_2$) δ[ppm] 7.37 – 7.30 (m, 10H), 7.20 – 7.13 (m, 6H), 6.68 (ddt, *J* = 2.1, 1.3, 0.7 Hz, 1H), 3.50 – 3.32 (m, 1H), 3.16 (ddd, *J* = 29.5, 7.8, 6.4 Hz, 1H), 2.90 – 2.66 (m, 2H), 1.77 – 1.59 (m, 1H), 1.44 – 1.37 (m, 19H), 1.36 – 1.28 (m, 2H), 0.94 – 0.77 (m, 6H). **$^{13}$C NMR** (126 MHz, $CD_2Cl_2$) δ[ppm] 174.80, 173.28, 143.16, 143.11, 138.72, 138.21, 137.81, 130.21, 128.38, 128.35, 128.28, 119.60, 80.81, 80.73, 75.47, 60.73, 60.54, 59.14, 59.07, 43.43, 43.28, 32.79, 32.67, 28.24, 25.26, 25.10, 23.00, 22.95, 22.57, 22.54. **IR** (v/cm$^{-1}$, neat): 3346.85, 2968.87, 2955.38, 2930.79, 2867.63, 1723.09, 1597.73, 1559.17, 1476.24, 1444.42, 1391.39, 1366.32, 1324.37, 1257.36, 1237.59, 1146.96, 1086.69, 1034.62, 1000.39. **HRMS** (ESI): calculated for $C_{39}H_{49}N_3O_4$ $[M+H]^+$: 624.3796, found: 624.3792.

***Characterization of compound 17:*** **$^{1}$H NMR** (400 MHz, $CD_2Cl_2$) δ[ppm] 7.25 – 7.23 (m, 1H), 6.91 – 6.89 (m, 1H), 4.75 (s, 2H). **$^{13}$C NMR** (100 MHz, $CD_2Cl_2$) δ[ppm] 119.33, 105.61, 105.24, 63.08. **$^{19}$F NMR** (377 MHz, $CD_2Cl_2$) δ[ppm] -68.04. **HRMS** (ESI): calculated for $C_6H_5ClFNO$ $[M+H]^+$: 162.0116, found: 162.0114.

***Characterization of compound 18.*** Compound **18** was obtained as a mixture of two diastereomers after column chromatography with a diastereomers mixture of (*S,S*):(*S,R*) 3:1 as judged per NMR. **$^{1}$H NMR** (400 MHz, $CDCl_3$) δ[ppm] 7.50 (s, 1H), 6.97 (s, 1H), 6.97 (s, 1H), 6.49 (s, 1H), 5.51 – 5.18 (m, 2H), 3.22 (dt, *J* = 48.8, 6.7 Hz, 2H), 2.84 – 2.69 (m, 2H), 1.42 (s, 18 H), 1.70 – 1.66 (m, 1H), 1.37 – 1.25 (m, 2H), 0.89 (dd, *J* = 8.3, 7.2 Hz, 6H). **$^{13}$C NMR** (100 MHz, $CDCl_3$) δ[ppm] 174.21, 172.25, 162.01, 138.08, 129.44, 125.44, 119.31, 105.83, 105.48, 60.44, 59.39, 46.95, 42.76, 28.38, 28.17, 25.01, 22.88, 22.24. **$^{19}$F NMR** (377 MHz, $CD_2Cl_2$) δ[ppm] -65.01. **IR** (v/cm$^{-1}$, neat): 2975.62, 2963.09, 2933.20, 2875.34, 2358.52, 2335.37, 1723.09, 1606.41, 1568.81, 1491.67, 1455.99, 1408.75, 1368.25, 1276.65, 1250.61, 1225.54, 1147.77, 1031.73. **HRMS** (ESI): calculated for $C_{26}H_{38}ClFN_4O_4$ $[M+H]^+$: 525.2638, found: 525.2645.

***Characterization of the (S,S)-F-Aza-MLN-4760*** · *TFA salt*: ***F-Aza-MLN-4760*** · TFA salt was obtained after HPLC purification as a pure (*S*,*S*) diastereomer, identified by NMR. **$^{1}$H NMR** (400 MHz, $CD_3OD$)

δ[ppm] 9.02 (s, 1H), 7.66 (s, 1H), 7.32 (s, 1H), 6.95 (s, 1H), 5.74 – 5.62 (m, 2H), 5.68 (dd, $J$ = 25.6, 17.2 Hz, 2H), 3.83 (t, $J$ = 5.3 Hz, 1H), 3.77 (dd, $J$ = 7.8, 6.2 Hz, 1H), 3.22 (ddd, $J$ = 41.8, 17.1, 6.4 Hz, 2H), 1.98-1.80 (m, 1H), 1.75 – 1.57 (m, 2H), 0.98 (d, $J$ = 6.5 Hz, 6H). **$^{13}$C NMR** (100 MHz, $CD_3OD$) δ[ppm] 173.49, 172.31, 136.39, 120.04, 119.62, 106.68, 106.30, 58.87, 40.74, 25.56, 24.49, 21.52, 20.94. **$^{19}$F NMR** (377 MHz, $CD_3OD$) δ[ppm] -67.53, -77.09. **IR** (v/cm$^{-1}$, neat): 3126.04, 3046.98, 2963.09, 2873.42, 1725.01, 1666.2, 1608.34, 1571.7, 1412.6, 1299.79, 1179.26, 1133.94. **HRMS** (ESI): calculated for $C_{18}H_{22}ClFN_4O_4$ [M+H]$^+$: 413.1386, found: 413.1382.

### 4. Chemical synthesis of the precursors for radiofluorination

#### 4.1. Synthesis of the trimethylstannane-based precursor 1 for the production of [$^{18}$F]F-MLN-4760

**Purpose**: A trimethylstannane-based radiofluorination precursor was designed and synthesized for the preparation of [$^{18}$F]F-MLN-4760.

**Methods:** The precursor molecule was synthesized following the synthetic route as described for the production of F-Aza-MLN-4760 with slight modifications (Scheme S3).

**Scheme S3.** Synthesis of the radiofluorination precursor **1**.

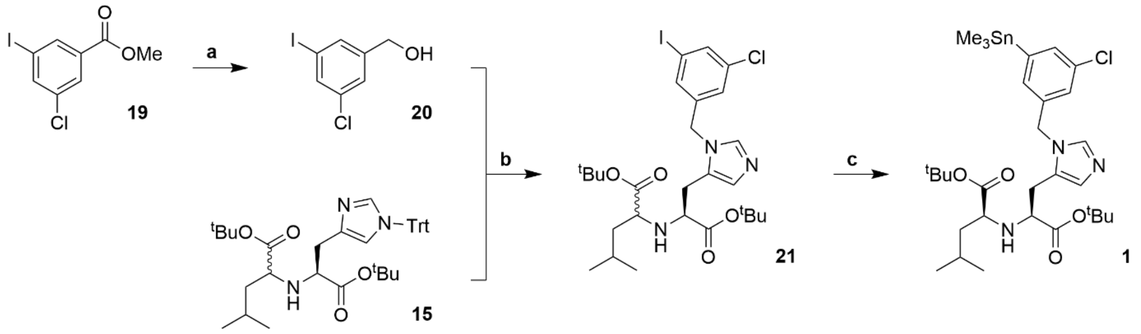


Reaction conditions: (**a**) $LiAlH_4$, THF, 0 °C to RT, 2 h; (**b**) $Tf_2O$, DIPEA, DCM, –78 °C to RT, overnight; (**c**) $Sn_2Me_6$, $Pd(PPh_3)_4$, THF, 100 °C, 45 min.

***Synthesis of intermediate* 20**. $LiAlH_4$ (2.05 g, 5.40 mmol, 0.80 equiv) was suspended in THF (18 mL), cooled to 0 °C, and a solution methyl 3-chloro-5-iodobenzonate (**19**, 2.0 g, 6.75 mmol, 1.00 equiv) in THF (6 mL) was added dropwise. The reaction mixture was stirred for 2 h at RT. The reaction was quenched with deionized $H_2O$ (1 mL), filtered over celite, and the resulting residue was washed with EtOAc. The filtrate was then dried over $Na_2SO_4$, filtered and concentrated under reduced pressure. The resulting yellow oil was purified by silica gel chromatography (hexane:EtOAc, 20:1 to 1:1 (*v/v*)) to obtain compound **20** as an off-white solid (1.19 g, 66% yield).

***Synthesis of intermediate 21***. DIPEA (117 µL, 675 µmol, 1.10 equiv) was added to a solution of compound **20** (180 mg, 675 µmol, 1.10 equiv) in DCM (560 µL) before cooling to –78 °C. A solution of trifluoromethanesulfonic anhydride (113 µL, 675 µmol, 1.10 equiv) in DCM (2 mL) was added under argon and the resulting mixture was stirred for 30 min at –78°C followed by slow addition of compound **15** (382 mg, 614 µmol, 1.00 equiv) in DCM (610 µL). The reaction mixture was stirred at RT overnight. The reaction mixture was washed with deionized $H_2O$ (2 x 5 mL) and brine (5 mL), dried over $Na_2SO_4$ and concentrated by evaporation under reduced pressure. The crude was purified by silica gel chromatography (hexane/EtOAc 4:1 (*v/v*), then EtOAc, then 5% MeOH in EtOAc) to afford a diastereomeric mixture of intermediate **21** (368 mg, 94% yield, (*S,S*):(*S,R*) 3:1).

***Synthesis of the aryl-trimethylstannane precursor 1***. To a solution of compound **21** (66.0 mg, 100 µmol, 1.00 equiv) and tetrakis(triphenylphosphine)palladium$^0$ ($Pd(PPh_3)_4$, 13.0 mg, 10.0 µmol, 0.10 equiv) in toluene (2 mL) was added dropwise hexamethylditin (924 µL, 400 µmol, 4.00 equiv). The resulting solution was stirred at 100 °C for 45 min before cooling to RT. The solvent was removed under reduced pressure and the crude product was purified by silica gel chromatography (hexane, then hexane/EtOAc 4:1 (*v/v*), then EtOAc). The collected fractions containing the compound of interest were further purified by RP-HPLC (35-95% MeCN in MilliQ $H_2O$ + 0.1% TFA). The desired (*S,S*)-diastereomer (precursor **1**) was identified based on the assumption that the sequence of the eluted diastereoisomers would be the same as that of the diastereoisomers of MLN-4760. Fractions containing the pure (*S,S*)-diastereomer of compound **1** were collected, combined and neutralized with aq. saturated $NaHCO_3$ to pH 7 before proceeding with the extraction of the product using DCM (2 x 10 mL). The combined organic layers were then dried over $Na_2SO_4$ and concentrated to afford the radiofluorination precursor **1** as a colorless oil (20 mg, 30% yield).

**Results:** The NMR and HRMS data of all synthesis intermediates and the final compound **1** are given below. The desired radiofluorination precursor **1** was obtained in an overall yield of 8%, calculated over the longest synthetic path which comprised 5 steps. The chemical purity of the final compound was determined to be >99% by LCMS and NMR.

***Characterization of compound 20:*** **$^1$H NMR** (500 MHz, $d_6$-DMSO) δ[ppm] 7.69 – 7.66 (m, 1H), 7.64 (dq, *J* = 2.8, 1.4 Hz, 1H), 7.38 (dq, *J* = 2.2, 1.1 Hz, 1H), 5.41 (qd, *J* = 5.1, 2.3 Hz, 1H), 4.47 (d, *J* = 5.5 Hz, 2H). **$^{13}$C NMR** (126 MHz, DMSO) δ[ppm] 147.24, 134.28, 133.73, 133.62, 125.66, 95.11, 61.36. **IR** (v/cm$^{-1}$, neat): 3314.07, 3212.34, 3065.3, 2933.68, 2867.63, 1581.83, 1558.2, 1425.14, 1358.12, 1310.39, 1206.26, 1102.6, 1078.01, 1013.41. **HRMS** (ESI): calculated for $C_7H_6ClIO$ [M]: 267.9144, found: 267.9146.

***Characterization of compound 21:*** After column chromatography, compound **21** was obtained as a mixture of two diastereomers in a (*S*,*S*)-to-(*S*,*R*) ratio of 3:1, identified by NMR. **$^1$H NMR** (500 MHz, $CD_2Cl_2$) δ[ppm] 8.67 (d, *J* = 32.5 Hz, 1H), 7.77 (dt, *J* = 3.4, 1.6 Hz, 1H), 7.49 (dt, *J* = 36.2, 1.5 Hz, 1H), 7.40 (d, *J* = 11.0 Hz, 1H), 7.18 (dt, *J* = 31.5, 1.7 Hz, 1H), 5.67 – 5.36 (m, 2H), 3.56 – 3.09 (m, 2H), 3.04 – 2.89 (m, 2H), 1.74 – 1.63 (m, 1H), 1.44 (dd, *J* = 13.3, 6.8 Hz, 21H), 0.91 (dd, *J* = 11.0, 6.6 Hz, 4H), 0.88 (d, *J* = 0.7 Hz, 1H), 0.83 (d, *J* = 6.6 Hz, 1H). **$^{13}$C NMR** (126 MHz, $CD_2Cl_2$) δ[ppm] 173.12,

170.95, 170.40, 138.02, 137.89, 136.79, 136.63, 136.04, 135.93, 135.39, 135.34, 134.86, 134.51, 130.98, 130.91, 127.19, 126.88, 119.97, 119.79, 117.30, 115.00, 94.93, 94.82, 83.27, 81.84, 59.49, 59.18, 59.06, 49.32, 49.22, 42.08, 41.88, 29.70, 27.75, 27.71, 27.66, 27.43, 26.71, 24.88, 22.33, 22.12, 22.08, 21.88. **IR** (v/cm$^{-1}$, neat): 3121.22, 3057.58, 2964.05, 2924.52, 2870.04, 2852.2, 2359.48, 2342.12, 1728.87, 1668.12, 1586.16, 1559.65, 1457.44, 1431.4, 1394.76, 1369.21, 1257.84, 1194.69, 1143.1, 1022.09. **HRMS** (ESI): calculated for $C_{27}H_{39}ClIN_3O_4$ $[M+H]^+$: 632.1747, found: 632.1749.

***Characterization of compound 1:*** Compound **1** was obtained after HPLC purification of the pure (*S*,*S*) diastereomer, identified by NMR. **$^1$H NMR** (500 MHz, $CD_2Cl_2$) δ[ppm] 7.44 – 7.37 (m, 2H), 7.14 (dd, *J* = 1.6, 0.8 Hz, 1H), 6.94 – 6.86 (m, 2H), 5.19 – 5.07 (m, 2H), 3.36 (t, *J* = 6.8 Hz, 1H), 3.17 (dd, *J* = 7.9, 6.3 Hz, 1H), 2.74 (dddd, *J* = 54.0, 15.1, 6.8, 0.8 Hz, 2H), 1.69 (ddd, *J* = 13.0, 7.5, 6.5 Hz, 1H), 1.41 (d, *J* = 9.5 Hz, 21H), 0.89 (dd, *J* = 9.7, 6.6 Hz, 6H), 0.29 (s, 9H). **$^{13}$C NMR** (126 MHz, $CD_2Cl_2$) δ[ppm] 174.12, 172.50, 146.04, 138.03, 137.74, 134.82, 134.76, 132.16, 128.34, 127.65, 126.38, 81.32, 80.70, 59.88, 58.95, 47.93, 42.69, 29.68, 28.31, 27.78, 27.72, 24.83, 22.48, 21.96, -9.75. **IR** (v/cm$^{-1}$, neat): 2956.34, 2923.56, 2870.04, 2852.2, 2362.37, 2335.37, 1725.98, 1558.68, 1488.78, 1457.44, 1435.26, 1392.35, 1366.8, 1341.73, 1250.62, 1212.04, 1150.81, 1109.35. **HRMS** (ESI): calculated for $C_{30}H_{48}ClN_3O_4Sn$ $[M+H]^+$: 670.2428, found: 670.2419. **$[\alpha]_{546}^{25}$**= −18.8000 (c=0.05, DCM).

#### 4.2. Synthesis of the pyridine-based precursor 2 for the production of [$^{18}$F]F-Aza-MLN-4760

**Purpose**: A pyridine-based precursor was designed and synthesized for the preparation of [$^{18}$F]F-Aza-MLN-4760.

**Methods:** The precursor molecule was synthesized following the same synthetic route as described for the production of F-Aza-MLN-4760 but, in this case, (2,6-dichloropyridin-4-yl)methanol was used in place of (2-chloro-6-fluoropyridin-4-yl)methanol (Scheme S4).

**Scheme S4** Synthesis scheme of the radiofluorination precursor **2**

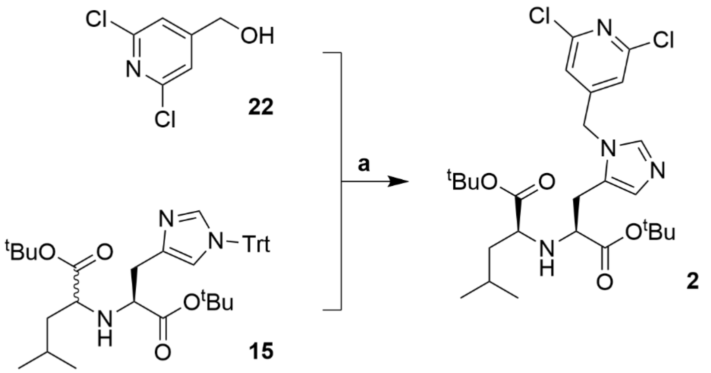


Reaction conditions: (**a**) $Tf_2O$, DIPEA, DCM, –78 °C to RT, overnight;

***Synthesis of the pyridine-based precursor (2).*** DIPEA (33 µL, 160 µmol, 1.20 equiv) was added to a solution of (2,6-dichloropyridin-4-yl)methanol (**22**, 34.2 mg, 190 µmol, 1.20 equiv) in DCM (1 mL) before cooling to –78 °C. A solution of trifluoromethanesulfonic anhydride (32.1 µL, 190 µmol, 1.20 equiv) in DCM (1 mL) was added under argon and the resulting mixture was stirred for 30 min at –78 °C followed by slow addition of compound **15** (100 mg, 160 µmol, 1.00 equiv) in DCM (500 µL). The reaction mixture was stirred at RT overnight. The reaction mixture was diluted with DCM (10 mL), washed with deionized $H_2O$ (2 x 5 mL) and brine (5 mL), dried over $Na_2SO_4$ and concentrated by evaporation under reduced pressure. The crude product was purified by silica gel chromatography (100% DCM, then DCM/MeOH 95:5 (*v/v*)). The collected fractions containing the compound of interest were further purified by RP-HPLC (10-90% MeCN in MilliQ $H_2O$ + 0.1% TFA). The desired (*S,S*)-diastereomer (precursor **2**) was identified based on the assumption that the sequence of the eluted diastereoisomers would be the same as that of the diastereoisomers of MLN-4760. Fractions containing the pure (*S,S*)-diastereomer of compound **2** were collected, combined and neutralized with aq. saturated $NaHCO_3$ to pH 7 before proceeding with the extraction of the product using DCM (2 x 10 mL). The combined organic layers were then dried over $Na_2SO_4$ and concentrated to afford the radiofluorination precursor **2** as a colorless oil (85 mg, 98% yield).

**Results:** In order to synthesize [$^{18}$F]F-Aza-MLN-4760, a pyridine-based precursor was designed and synthesized. The NMR and HRMS data of all synthesis intermediates and the final compound **2** are given below. The desired radiofluorination precursor **2** was obtained in an overall moderate yield (28%) calculated over the longest synthetic path (4 steps). The chemical purity of the final compound was determined to be >99% by LCMS and NMR.

***Characterization of compound 2:*** Compound **2** was obtained after HPLC purification as a pure (*S,S*) diastereomer, identified based on the NMR. **$^1$H NMR** (400 MHz, $CD_3OD$) δ[ppm] 7.79 (s, 1H), 7.10 (s, 2H), 6.92 (s, 1H), 5.43 – 5.33 (m, 2H), 3.28 (dd, *J* = 8.3, 5.9 Hz, 1H), 3.18 (t, *J* = 7.5 Hz, 1H), 2.80 (ddd, *J* = 42.4, 15.0, 8.2 Hz, 2H), 1.85-1.63 (m, 1H), 1.49– 1.26 (m, 20H), 0.92 (dd, *J* = 10.2, 6.8 Hz, 2H). **$^{13}$C NMR** (377 MHz, $CD_3OD$) δ[ppm] 173.98, 172.62, 153.18, 153.15, 138.30, 128.07, 127.43, 120.65, 81.65, 81.03, 59.80, 58.44, 45.89, 42.35, 27.10, 26.90, 24.60, 21.59, 21.25. **IR** (v/cm$^{-1}$, neat): 2957.3, 2931.27, 2866.67, 2361.41, 1724.05, 1586.16, 1547.59, 1488.78, 1453.1, 1383.68, 1367.28, 1336.43, 1249.65, 1215.9, 1149.37, 1108.87. **HRMS** (ESI): calculated for $C_{26}H_{38}Cl_2N_4O_4$ $[M+H]^+$: 541.2343, found: 541.2333. **$[\alpha]_{546}^{25}$**= − 14.0000 (c=0.05, DCM).

### 5. Radiosynthesis

#### 5.1 Radiolabeling of [$^{18}$F]F-MLN-4760

**Purpose:** The $^{18}$F-based radiotracer was synthesized by $^{18}$F-fluorination of precursor **1**.

**Methods:** The radiofluorination was carried out using the aryl-trimethylstannane precursor (compound **1**). [$^{18}$F]Fluoride was produced by the bombardment of the 98% enriched $^{18}$O-water target via the

$^{18}O(p,n)^{18}F$ nuclear reaction using a medical cyclotron (18-MeV, IBA, Belgium) installed at ETH Zurich. The aqueous solution was transferred from the cyclotron to the hot-cell and the [$^{18}$F]Fluoride was trapped on an anion-exchanger cartridge (Sep-Pak Accell Plus QMA Plus Light Cartridge, WAT023525, preconditioned with 5 mL EtOH, 5 mL potassium triflate in MilliQ $H_2O$ (90 mg/mL), and 5 mL MilliQ $H_2O$). A mixture of potassium triflate (450 µL, 10 mg/mL in MilliQ $H_2O$), $K_2CO_3$ (130 µL, 1 mg/mL in MilliQ $H_2O$) and MeCN (500 µL) was applied to elute the activity to the reaction vial. After azeotropic drying with MeCN (2 x 1.0 mL), the reaction vial was purged with air. The precursor **1** (4.0-5.0 mg) in 300 µL of dimethylacetamide (DMA) and 16-17 mg $Cu(OTf)_2(Py)_4$ in 300 µL DMA were added sequentially to the residue and the reaction mixture was heated at 110 °C for 10 min. The deprotection was carried out by adding 600 µL orthophosphoric acid (85% (*m*/*m*)) and subsequent stirring at 110 °C for 15 min. After dilution with phosphate buffered saline (PBS, 1.0 mL), the mixture was processed using semi-preparative HPLC purification (Phenomenex Luna, 10 µm, C18, 100 Å, 250 x 10 mm, mobile phase A: 0.1% $H_3PO_4$ in MilliQ $H_2O$, mobile phase B: MeCN, gradient method: 0.0−5.0 min, 5% B; 5.0−35.0 min, 5−25% B; and 35.0−40.0 min, 25−95% B; flow =4 mL/min; λ=254 nm). The collected fraction was diluted with 20 mL MilliQ $H_2O$ and passed through a C18 light cartridge (Waters, WAT023501, preconditioned with 5 mL EtOH and 5 mL MilliQ $H_2O$). After washing the cartridge with MilliQ $H_2O$ (5 mL), **[$^{18}$F]F-MLM-4760** was eluted with 0.5 mL EtOH. The final product was formulated to obtain a solution containing 10% (*v*/*v*) EtOH in PBS. The radioactive products were analyzed with an Agilent 1100 series HPLC system, equipped with a UV detector, and a GabiStar radiodetector (Raytest) using Phenomenex column (Gemini® 5 µm C18 110 Å, 150 x 4.6 mm, mobile phase A: 0.01% $H_3PO_4$ in MilliQ $H_2O$, mobile phase B: MeCN, gradient method: 0.0−12.0 min, 5−30% B; 12−15 min, 30-50% B; 15.0−18.0 min, 5% B with the flow of 4 mL/min at 254 nm). The reactions scheme is reported in the main article (Scheme 1).

**Results: [$^{18}$F]F-MLN-4760** was obtained in a radiochemical purity of >99% with a radiochemical yield of up to 6.4%. Molar activities ranged from 21 to 38 GBq/µmol (n = 5) at the end of the synthesis. The chemical identity and stereocenter configuration of the final product were confirmed by HPLC co-injection of (*S,S*)-F-MLN-4760 (Fig. S2).

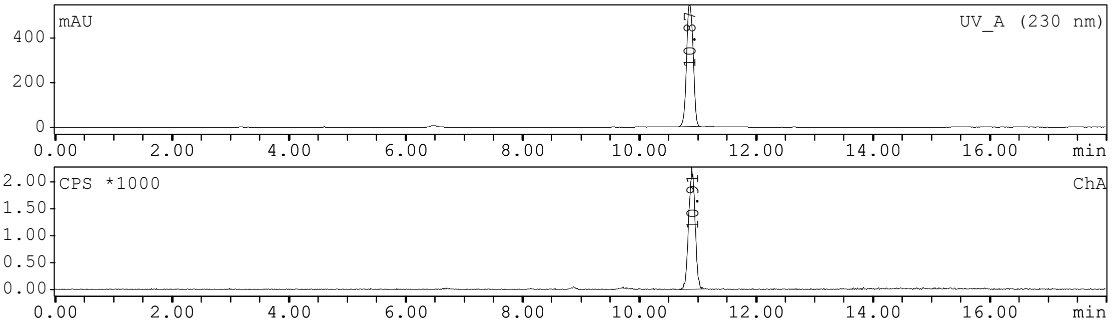


**Fig. S2** Chromatograms obtained from co-injecting the reference compound (*S,S*)-F-MLN-4760 and [$^{18}$F]F-MLN-4760

#### 5.2 Radiolabeling of [$^{18}$F]F-Aza-MLN-4760

**Purpose:** [$^{18}$F]F-Aza-MLN-4760 was synthesized by $^{18}$F-fluorination of precursor **2**, followed by deprotection and HPLC purification.

**Methods:** The radiofluorination was carried out using the pyridine-based precursor **2**. [$^{18}$F]Fluoride was produced as described in Section 5.1. The aqueous solution was transferred from the cyclotron to the hot-cell and the [$^{18}$F]Fluoride was trapped on an anion-exchanger cartridge (Sep-Pak Accell Plus QMA Plus Light Cartridge, WAT023525, preconditioned with 10 mL of $K_2CO_3$ 0.5 M and 10 mL MilliQ $H_2O$). A mixture of $Cs_2CO_3$ (175 µL, 16 mg/mL in MilliQ $H_2O$), Kryptofix® 222 (175 µL, 37 mg/mL in MilliQ $H_2O$) and MeCN (300 µL) was applied to elute the activity to the reaction vial. After azeotropic drying with MeCN (2 x 1.0 mL), the reaction vial was purged with air. The precursor **2** (3.5-4.5 mg) in 500 µL of dimethylsulfoxide (DMSO) was added to the residue and the reaction mixture was heated at 195 °C for 20 min. The deprotection was carried out by adding HCl (1.0 mL, 4 M) and subsequent stirring at 80 °C for 20 min. After neutralization performed by addition of NaOH (1.0 mL, 4 M) and $Na_2HPO_4$ (500 µL, 75 mg/mL), the mixture was processed using semi-preparative HPLC purification (Phenomenex Luna, 10 µm, C18, 100 Å, 250 x 10 mm; mobile phase A: PBS pH= 7.4, mobile phase B: MeCN, gradient: 0.0−5.0 min, 5% B; 5.0−35.0 min, 5−15% B and 35.0−40.0 min, 15−95% B with a flow rate of 4 mL/min; λ=254 nm). The collected fraction was acidified with 10 mL HCl (0.5 M) and passed through a cations-exchange cartridge (Oasis MCX 30 mg, 1cc, preconditioned with 10 mL EtOH and 10 mL MilliQ $H_2O$). After rinsing the cartridge with MilliQ $H_2O$ (5 mL), **[$^{18}$F]F-Aza-MLN-4760** was eluted with 2.0 mL of 10% (*v/v*) EtOH in PBS pH 7.4. The radioactive products were analyzed using the same HPLC system as described in Section 3.1.

**Results: [$^{18}$F]F-Aza-MLN-4760** was obtained in a radiochemical purity of >99% with a radiochemical yield of up to 1.5%. Molar activities ranged from 78 to 81 GBq/µmol (n = 3) at the end of the synthesis. The chemical identity and stereocenter configuration of the final product were confirmed by HPLC co-injection of the (*S,S*)-F-Aza-MLN-4760 (Fig. S3). The radiolytic stability of [$^{18}$F]F-Aza-MLN-4760 was investigated over 2 h using HPLC. The results confirmed that the radiotracer remained stable over 2 h in the formulated solution.

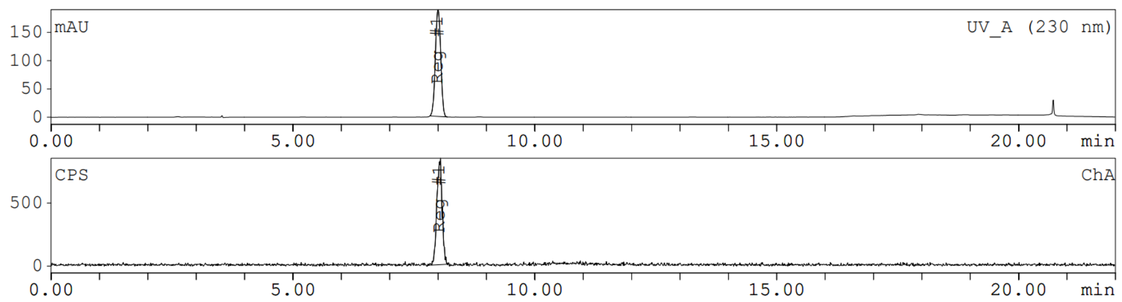


**Fig. S3** HPLC-chromatograms obtained from co-injecting the reference compound (*S,S*)-F-Aza-MLN-4760 and [$^{18}$F]F-Aza-MLN-4760

### 6. In Vitro stability of [$^{18}$F]F-MLN-4760

**Purpose:** The radiolytic stability of [$^{18}$F]F-MLN-4760 and [$^{18}$F]F-MLN-4760 as well as their stability in human and murine blood plasma was investigated.

**Method:** The radiolytic stability of [$^{18}$F]F-MLN-4760 and [$^{18}$F]F-Aza-MLN-4760 formulated in PBS pH 7.4 containing 10% EtOH was investigated over 3 h incubation at RT. Quality control was performed using a Merck Hitachi LaChrom HPLC system equipped with a D-7000 interface, a L-7200 autosampler, a radioactivity detector (LB 506 B; Berthold) and a L-7100 pump connected with a C18 column (Xterra$^{TM}$, 5 µm, 4.6×150 mm, Waters, Milford, MA, U.S.A.). The radiotracers were eluted using a linear gradient (5-80% MeCN in MilliQ $H_2O$ with 0.1% TFA) over 15 min at a flow rate of 1.0 mL/min.

[$^{18}$F]F-MLN-4760 and [$^{18}$F]F-Aza-MLN-4760 were incubated for up to 3 h in murine blood plasma (Lot: 32321, Rockland Inc.), human blood plasma (Blood donation SRK Aargau-Solothurn, Switzerland) or in 0.9% NaCl as a control (~10 MBq/200 µL) at 37 °C in a shaker. Thin layer chromatography was performed with an aliquot of the blood plasma samples after 1 h and 3 h using TLC reversed phase C-18 plates (TLC silica gel 60 RP-18; Merck). A mixture of 10% ammonium acetate in MilliQ $H_2O$ (50%, *v/v*) and MeOH (50%, *v/v*) was employed as mobile phase. Under these conditions, the intact radiotracer stayed at the starting line while potential smaller fragments or free [$^{18}$F]Fluoride migrated. A second chromatographic separation was performed using the same TLC reversed phase C-18 plates as a stationary phase but a mixture of citrate buffer pH 5.5 (60%, *v/v*) and MeCN (40%, *v/v*) as the mobile phase. In this case, the radiotracer migrated while potential smaller fragments or free [$^{18}$F]Fluoride stayed at the baseline. The TLC plates were analyzed using a storage phosphor system (Cyclone Plus, Perkin Elmer). The quantification of the signals was performed using OptiQuant software (version 5.0, Bright Instrument Co Ltd., Perkin Elmer$^{TM}$). The chromatograms were analyzed by determination of the peak area of the radiotracer as well as fragments of unknown structure or released [$^{18}$F]Fluoride. The quantity of the intact product was expressed as percentage of the sum of integrated peak areas of the entire chromatogram.

**Results:** HPLC analysis of samples of [$^{18}$F]F-MLN-4760 and [$^{18}$F]F-Aza-MLN-4760 showed that both radiotracers were radiolytically stable (> 97% intact radiotracer) for up to 3 h in the formulated solution (Fig. S4).

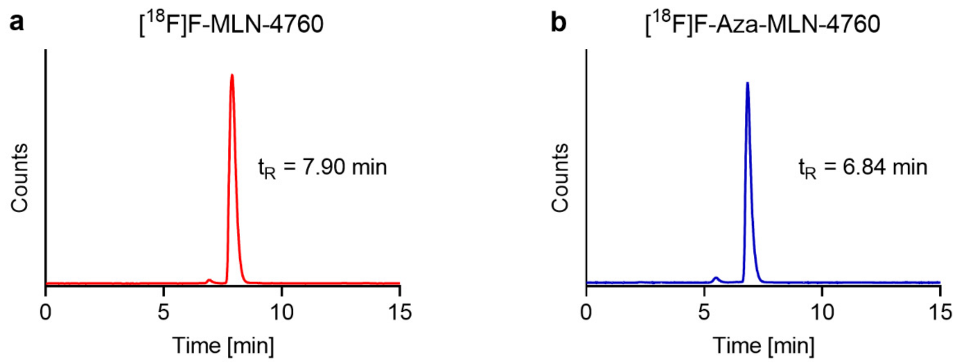


**Fig. S4** Chromatograms of [$^{18}$F]F-MLN-4760 (**a**) and [$^{18}$F]F-Aza-MLN-4760 (**b**) incubated for 3 h in their formulated solutions

No degradation products of [$^{18}$F]F-MLN-4760 or [$^{18}$F]F-Aza-MLN-4760 were observed for up to 3 h incubation in mouse (> 99% and > 97%, respectively) and human blood plasma (> 99% and > 97%, respectively).

#### 7. *n*-Octanol/PBS distribution coefficient

**Purpose:** The *n*-octanol/PBS distribution coefficient (logD value) was determined to assess the hydrophilic/lipophilic character of the $^{18}$F-based radiotracers.

**Methods:** A fixed amount of the respective radiotracers (1 MBq, 25 µL) was added to radio-immunoassay (RIA) tubes containing a mixture of PBS pH 7.4 (1475 µL) and *n*-octanol (1500 µL). After vortexing vigorously for 1 min, the tubes were centrifuged (560 rcf; 6 min) to obtain phase separation. The quantity of activity in the organic and aqueous phases was determined in a γ-counter (Perkin Elmer, Wallac Wizard 1480). The distribution coefficients were expressed as the logarithm of the ratio of counts per minute (cpm) measured in the *n*-octanol phase to the cpm measured in the PBS pH 7.4 phase and indicated as the average of three independent measurements (± standard deviation, SD), each performed with five replicates.

**Results:** The result is reported in the main article.

#### 8. Cell culture

HEK-ACE2 and HEK-ACE, respectively, were custom-made by Innoprot (Innovative Technologies in Biological Systems S.L. Bizkaia, Spain) using HEK-293 cells that were transfected with hACE2 and human ACE, respectively. They were cultured in Dulbecco's Modified Eagle Medium (DMEM) supplemented with non-essential amino acids, fetal calf serum and antibiotics. Hygromycin B was added to maintain the expression of ACE2 and ACE, respectively. The HEK cells were cultured under standard conditions at 37 °C and 5% $CO_2$ and subcultured using PBS/EDTA and trypsin.

### 9. ACE2-binding affinity of F-MLN-4760 and F-Aza-MLN-4760

**Purpose:** The ACE2-binding affinity ($IC_{50}$ values) of F-MLN-4760, F-Aza-MLN-4760 and MLN-4760 was investigated using [$^{3}$H]MLN-4760 and HEK-ACE2 cells.

**Methods:** HEK-ACE2 cells (0.25 Mio in 0.5 mL culture medium) were seeded in poly-D-lysine-coated 48-well plates allowing cell adhesion and growth overnight. [$^{3}$H]MLN-4760 (RC Tritec AG, Teufen, Switzerland) was used as a radiotracer (10 µL, 3.1 pmol per well, 6.2 nM). Displacement of [$^{3}$H]MLN-4760 by increasing concentrations (100-0.01 µM) of MLN-4760, F-MLN-4760 or F-Aza-MLN-4760 was investigated. After a 1 h incubation time at 37 °C, the cells were rinsed with PBS (pH 7.4, 1 mL) followed by lysis using NaOH (1 M, 600 µL) and transfered to scintillation vials. After addition of 5 mL scintillation cocktail (Ultima Gold, Perkin Elmer®), the vials were vortexed and swirled in EtOH to reduce the static load before counting in a liquid scintillation analyzer (TRI-CARB® 2250CA, Packard). The counts were expressed as percentage of maximum uptake of [$^{3}$H]MLN-4760 plotted against the logarithmic concentration of the test agent (MLN-4760, F-MLN-4760 or F-Aza-MLN-4760) to obtain their $IC_{50}$ values using GraphPad Prism software (version 8.3.1). Two batches of MLN-4760, F-MLN-4760 and F-Aza-MLN-4760, respectively, were weighed and used for the displacement experiments, resulting in four independent experiments for each compound. Displacement curves were generated from the average values obtained from the four experiments, which yielded the $IC_{50}$ value indicated as the value and 95% confidence interval. The relative binding affinities were expressed as the reversed $IC_{50}$ values which were standardized to MLN-4760 set as 1.0. The datasets were analyzed for significance with the extra sum-of-squares F test using GraphPad Prism Software.

**Results:** The displacement curves of MLN-4760, F-MLN-4760 and F-Aza-MLN-4760 revealed the strongest binding for MLN-4760 ($IC_{50}$: 52 nM, 95% CI: 40-69 nM), followed by F-MLN-4760 ($IC_{50}$: 150 nM; 95% CI: 113-198 nM) and F-Aza-MLN-4760 ($IC_{50}$: 387 nM; 95% CI: 247- 607) (Fig. S5). The relative binding affinities are reported in the main article.

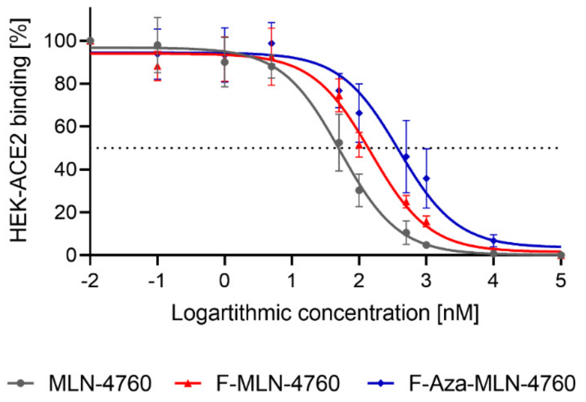


**Fig. S5** Displacement curve of [$^{3}$H]H-MLN-4760 obtained from an experiment performed on HEK-ACE2 cells using MLN-4760, F-MLN-4760 and F-Aza-MLN-4760.

### 10. Uptake of [$^{18}$F]F-MLN-4760 and [$^{18}$F]F-Aza-MLN-4760 in HEK-ACE2 and HEK-ACE cells

**Purpose:** The uptake of the radiotracers was investigated using HEK-ACE2 and HEK-ACE cells.

**Methods:** Uptake and internalization of [$^{18}$F]F-MLN-4760 and [$^{18}$F]F-Aza-MLN-4760 were determined using HEK-ACE2 and HEK-ACE cells according to the procedures previously established protocol [10]. The HEK-ACE2 and HEK-ACE cells were seeded in poly-D-lysine-coated 12-well plates, allowing cell adhesion and growth overnight. The cells were rinsed with PBS followed by the addition of [$^{18}$F]F-MLN-4760 or [$^{18}$F]F-Aza-MLN-4760 (25 µL, 0.2 MBq per well). In some cell samples, the radiotracer was co-incubated with an excess of MLN-4760 (2 µM) to block ACE2. After incubation of the cells for 1 h or 3 h, they were rinsed with only PBS or additionally with acidic stripping buffer (glycine buffer with NaCl 0.9%, pH 2.8) to determine the total uptake and internalization, respectively, of [$^{18}$F]F-MLN-4760 and [$^{18}$F]F-Aza-MLN-4760. Cell samples were lysed using NaOH (1 M, 1 mL) and transferred in RIA tubes. The cell samples were counted for activity using a γ-counter (Perkin Elmer, Wallac Wizard 1480). The results were expressed as a percentage of total added activity and normalized to an average content of ~0.3 mg protein per well.

**Results:** The HEK-ACE2 cell uptake of [$^{18}$F]F-MLN-4760 was 49 ± 10% and 67 ± 9% after 1 h and 3 h incubation, respectively, while considerably lower values were seen for [$^{18}$F]F-Aza-MLN-4760 (28 ± 10% and 37 ± 8%, after 1 h and 3 h, respectively) (Fig. S6a). Both radiotracers showed only moderate internalization (< 11%) after 3 h of incubation (Fig. S6b). Co-incubation of the HEK-ACE2 cells with an excess of MLN-4760 blocked ACE2 and prevented the uptake of the radiotracers almost completely (<1.5%, Fig. S6c). The uptake in HEK-ACE cells is reported in the main article.

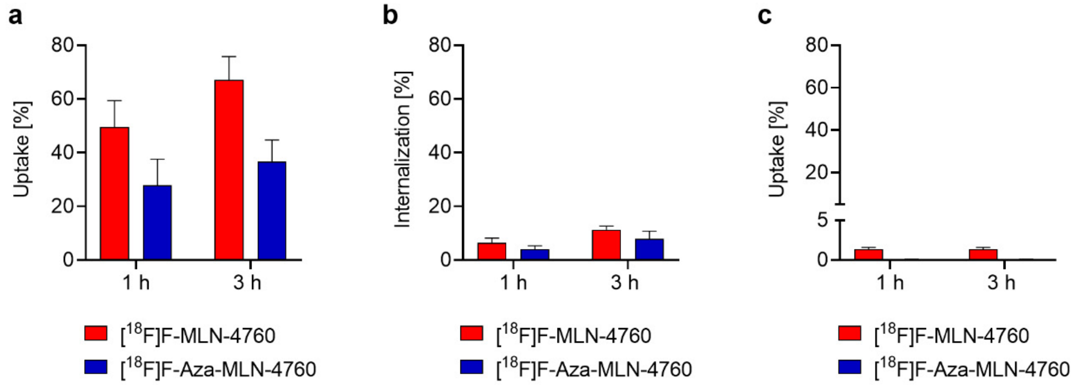


**Fig. S6** (**a/b**) Uptake and internalization of [$^{18}$F]F-MLN-4760 and [$^{18}$F]F-Aza-MLN-4760 in HEK-ACE2 cells after 1 h and 3 h incubation time. (**c**) Uptake of [$^{18}$F]F-MLN-4760 and [$^{18}$F]F-Aza-MLN-4760 co-incubated with an excess of MLN-4760 to block ACE2

### 11. Autoradiography on murine tissue sections

**Purpose**: In vitro autoradiography studies were performed on frozen tissue sections of the heart, lung, kidney and brain tissue from CD1 nude mice. The aim was to investigate the specificity of the radiotracers binding on mouse tissue.

**Methods**: Frozen tissue sections (10 µm) from lungs, kidneys, heart, brain and HEK-ACE2 and HEK-ACE xenografts were collected from CD1 nude mice, embedded in Tissue-Tek® O.C.T. and frozen at -80°C. Sections of 10 µm thickness were prepared using a cryostat (Epredia™ Cryostar™ NX70 Cryostat, Microm International GmbH, Dreieich, Germany) on slides (Superfrost™, Plus Adhesion Microscope Slides, epredia). The sections were incubated Tris-HCl buffer (170 nM, pH 7.6, with 5 mM $MgCl_2$) with 0.25% BSA for 10 min before being exposed to [$^{18}$F]F-MLN-4760 or [$^{18}$F]F-Aza-MLN-4760 (225 kBq/ 150 µL) in Tris-HCl buffer with 1% (*w/v*) BSA for 60 min at RT. MLN-4760 (10 µM) was added to block ACE2 binding. After incubation, the tissue sections were rinsed twice for 5 min with Tris-HCl buffer containing BSA followed by rinsing the sections with Tris-HCl buffer without BSA and MiliQ $H_2O$. After drying the sections at RT, images were obtained using a storage phosphor imager and quantified using OptiQuant software (version 5.0). The tissue sections were exposed together with 1-3 µL of dilutions of the radiotracer solution of known activities (30-4500 Bq). The amount of activity bound to the target structure on the tissue sections was quantified by converting the signal intensity measured as digital light unit (DLU)/mm$^2$ to activity per area in Bq/mm$^2$. The specific ACE2 binding of the radiotracers on the xenografts and organ tissue section was obtained by subtracting the areas of tissue sections that were incubated with an excess of MLN-4760 from their adjacent sections (total activity per area).

**Results**: Specific binding of [$^{18}$F]F-MLN-4760 was observed in kidney, heart and lung tissue of mice expressing murine ACE2, however, [$^{18}$F]F-Aza-MLN-4760 revealed a reduced binding when incubated on the same organs resulting in close to no specific signal when incubated on kidney, lung and brain tissue (Fig. S7). The results from the activity quantification of the autoradiography are reported in the main article.

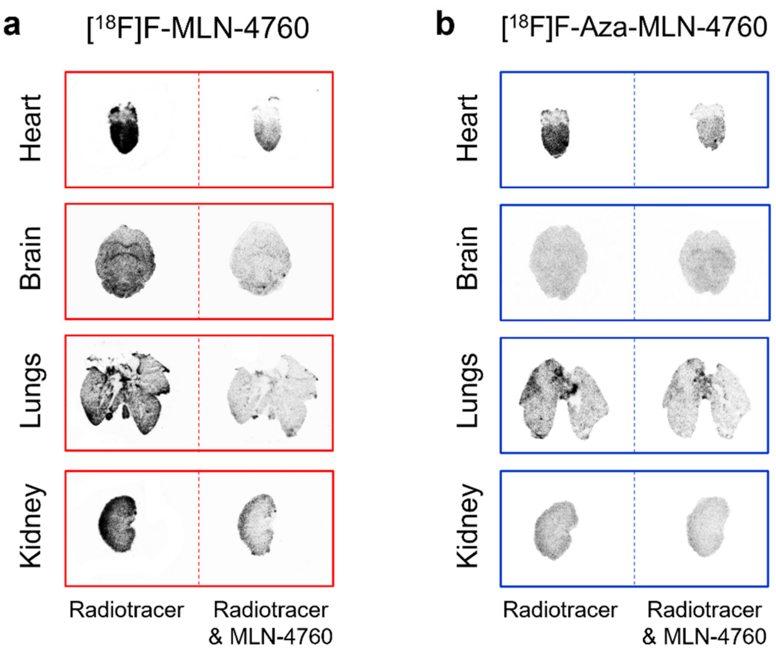


**Fig. S7 a/b** Representative autoradiograms obtained with [$^{18}$F]F-MLN-4760 and [$^{18}$F]F-Aza-MLN-4760 on frozen tissue sections of mice. **a** Autoradiograms of heart, brain, lung and kidney tissue incubated with [$^{18}$F]F-MLN-4760 only or in the presence of excess MLN-4760. **b** Autoradiograms of heart, brain, lung and kidney tissue incubated with [$^{18}$F]F-Aza-MLN-4760 only or in the presence of excess MLN-4760

### 12. Biodistribution studies of [$^{18}$F]F-MLN-4760 and [$^{18}$F]F-Aza-MLN-4760 in mice

**Purpose**: Biodistribution studies of [$^{18}$F]F-MLN-4760 were performed using HEK-ACE2 and HEK-ACE xenograft-bearing mice.

**Methods**: CD1 nude (Crl:CD1-*Foxn*$^{nu}$) mice were subcutaneously inoculated with HEK-ACE2 cells (6–8 × $10^6$ cells in 100 µL PBS) on the right shoulder and approximately one week later the same mice were inoculated with HEK-ACE (4 × $10^6$ cells in 100 µL PBS) on the left shoulder. After the formation of xenografts (100-300 mm$^3$), the mice were intravenously injected with [$^{18}$F]F-MLN-4760 or [$^{18}$F]F-Aza-MLN-4760 (5 MBq, 100 µL) diluted in NaCl 0.9% containing 0.05% bovine serum albumin (BSA). The EtOH content of the injection solutions was below 5%. The mice were sacrificed and dissected 15 min, 1 h or 3 h post injection (p.i.) of [$^{18}$F]F-MLN-4760 and 1 h p.i. of [$^{18}$F]F-Aza-MLN-4760. The results were listed as a percentage of the injected activity per gram of tissue mass (% IA/g), using counts of a defined volume of the original injection solution measured at the same time, resulting in decay-corrected values.

**Results**: The data of the biodistribution studies are presented and discussed in the main article and the values listed in Table S1.

**Table S1** Biodistribution data obtained 15 min, 1 h and 3 h after injection of HEK-ACE2 and HEK-ACE-bearing mice with [$^{18}$F]F-MLN-4760 and [$^{18}$F]F-Aza-MLN-4760. Decay-corrected data of accumulated activity are shown as % IA/g tissue, representing the average ± SD (n = 3)

| | **[$^{18}$F]F-MLN-4760** | | | **[$^{18}$F]F-Aza-MLN-4760** |
|---|---|---|---|---|
| | **15 min p.i.** | **1 h p.i.** | **3 h p.i.** | **1 h p.i.** |
| Blood | 0.70 ± 0.17 | 0.04 ± 0.01 | < 0.03 | 0.08 ± 0.02 |
| Heart | 0.47 ± 0.13 | 0.03 ± 0.01 | < 0.03 | 0.05 ± 0.01 |
| Lung | 0.63 ± 0.18 | 0.10 ± 0.06 | < 0.03 | 0.11 ± 0.02 |
| Spleen | 0.38 ± 0.09 | 0.06 ± 0.02 | < 0.03 | 0.14 ± 0.06 |
| Kidneys | 44 ± 8 | 5.1 ± 1.5 | 0.25 ± 0.02 | 2.2 ± 0.6 |
| Stomach | 2.2 ± 1.8 | 0.47 ± 0.44 | 0.07 ± 0.1 | 0.09 ± 0.03 |
| Intestines | 10 ± 13 | 28 ± 5 | 5.0 ± 4.8 | 10 ± 5 |
| Liver | 5.7 ± 3.3 | 1.6 ± 1.1 | 0.96 ± 1.2 | 0.84 ± 0.57 |
| Muscle | 0.23 ± 0.11 | 0.06 ± 0.01 | < 0.03 | 0.14 ± 0.19 |
| Bone | 0.22 ± 0.37 | 0.08 ± 0.02 | 0.07 ± 0.03 | 0.12 ± 0.03 |
| HEK-ACE2 xenograft | 11 ± 1 | 13 ± 2 | 5.8 ± 0.9 | 15 ± 2 |
| HEK-ACE xenograft | 0.26 ± 0.00 | 0.26 ± 0.16 | 0.07 ± 0.05 | 0.26 ± 0.00 |
| Salivary glands | 0.23 ± 0.12 | 0.04 ± 0.01 | < 0.03 | 0.07 ± 0.02 |
| Brain | 0.05 ± 0.00 | < 0.03 | < 0.03 | < 0.03 |

#### 13. PET/CT imaging studies

**Purpose**: PET/CT imaging studies were performed to evaluate the distribution profile of the radiotracers in mice bearing HEK-ACE2 and HEK-ACE xenografts.

**Methods**: PET/CT scans of xenograft-bearing mice were performed using a small-animal PET/CT scanner (G8, Perkin Elmer, U.S.A) as previously reported [10]. During PET/CT acquisitions, the mice were anesthetized using a mixture of isoflurane (1.5 – 2.0%) and oxygen. Static whole body PET scans of 10 min duration were performed at 15 min, 1 h and 3 h after intravenous injection of [$^{18}$F]F-MLN-4760 or [$^{18}$F]F-Aza-MLN-4760 (5 MBq, ~0.2 nmol, 100-200 μL) diluted in NaCl 0.9% containing 0.05% BSA. The PET scan was followed by a CT scan of 1.5 min duration. The acquisition of the data and their reconstruction was performed using the G8 PET/CT scanner software (version 2.0.0.10). The images were prepared using using VivoQuant post-processing software (version 3.5, inviCRO Imaging Services and Software, U.S.A.).

**Results**: The results are reported and discussed in the main article.
